# The Psychometrics of Automatic Speech Recognition

**DOI:** 10.1101/2021.04.19.440438

**Authors:** Lotte Weerts, Stuart Rosen, Claudia Clopath, Dan F. M. Goodman

## Abstract

Deep neural networks have had considerable success in neuroscience as models of the visual system, and recent work has suggested this may also extend to the auditory system. We tested the behaviour of a range of state of the art deep learning-based automatic speech recognition systems on a wide collection of manipulated sounds used in standard human psychometric experiments. While some systems showed qualitative agreement with humans in certain tests, in others all tested systems diverged markedly from humans. In particular, all systems used spectral invariance, temporal fine structure and speech periodicity differently from humans. We conclude that despite some promising results, none of the tested automatic speech recognition systems can yet act as a strong proxy for human speech recognition. However, we note that the more recent systems with better performance also tend to better match human results, suggesting that continued cross-fertilisation of ideas between human and automatic speech recognition may be fruitful. Our open source toolbox allows researchers to assess future automatic speech recognition systems or add additional psychoacoustic measures.

## 1 Introduction

It has been argued that deep neural networks are an excellent model of the visual system (Lindsay, 2021), in part thanks to their ability to predict neural responses based purely on training on behavioural tasks (Yamins et al., 2014). The impressive results of these modelling efforts (even if not universal; Jacob et al. 2021) have led to the suggestion that this approach should be extended more widely in neuroscience (Kriegeskorte, 2015; Richards et al., 2019). With recent advances in deep learning-based automatic speech recognition (ASR), there has therefore been an increased interest in comparing human speech recognition (HSR) with ASR (Rader et al., 2015; Stolcke and Droppo, 2017; Spille and Meyer, 2017; Kell et al., 2018; Spille et al., 2018; Hu et al., 2020; Kollmeier et al., 2020). ASR models may provide an opportunity to support or generate new hypotheses about the functioning of HSR. For instance, a recent comparison of ASR and HSR suggests ASR systems may be used to predict speech intelligibility for normal-hearing humans (Spille and Meyer, 2017). By applying explainable machine learning techniques, the use of ‘dip listening’ to predict speech under additive modulated maskers was identified in the ASR model. In another example, Kell et al. (2018) showed that a task-optimized neural network was able to replicate human auditory behaviour without being explicitly trained to do so. This model was subsequently used to accurately predict brain responses.

For an ASR system to be used as a model of human hearing, it is imperative that the system behaves as similarly as possible to HSR. As speech is inherently redundant, the same overall recognition performance may be achieved using different strategies and auditory cues. To test whether the same or different strategies and cues are being used, we systematically compared three publicly available ASR systems with human performance on a range of auditory dimensions. The test battery incorporates psychometric experiments that have been designed to measure the importance of a range of auditory cues as well as mechanisms thought to underlie human hearing, such as the use of temporal fine structure cues and dip listening. We have made the test battery freely available as part of HumanlikeHearing, a new open source Python toolbox we developed to allow researchers to directly compare HSR with ASR systems (https://github.com/neural-reckoning/HumanlikeHearing).

The results of our comparison suggest that on certain measures, such as sensitivity to clipping distortions, the performance of some of the tested ASR systems is comparable to that of HSR. In other cases, qualitative similarities were found for some ASR systems, such as sensitivity to spectral and temporal modulations, dip listening and masker periodicity. However, significant differences with human performance were found with respect to spectral invariance, the use of temporal fine structure and the role of periodicity in target speech for all ASR systems. Together, these results show that although some of the ASR systems are surprisingly robust to distortions without being trained on distorted data, there are still stark differences in the mechanisms employed and the relative importance of particular auditory cues compared to HSR. By sharing our test battery in the HumanlikeHearing toolbox, other researchers can now quickly and easily evaluate whether or not candidate ASR systems are a suitable proxy for HSR.

## 2 Results

We tested three freely available ASR models, each of which uses a different approach to model the temporal aspects of speech.

1. The Kaldi nnet3 chain model from the Kaldi Active Grammar project (Zurow et al., 2021) uses a hybrid Deep Neural Network and Hidden Markov Model (**DNN-HMM**; Povey et al. 2018). It takes as input Mel-frequency cepstral coefficient (MFCC) features, a classic set of features inspired by the human auditory system in which time is divided into short, overlapping ‘frames’, and the spectral envelope of the sound is computed for each frame. At each layer of the network, several frames of output from the previous layer are provided as input to give temporal context.
2. Mozilla’s DeepSpeech model (Mozilla, 2020) is a Long-Short-Term-Memory (**LSTM**) neural network, a widely used type of neural network that is designed to model long temporal relationships. It is a variant of the original DeepSpeech model that was introduced by Hannun et al. (2014). Like the DNN-HMM model, it uses MFCC features as its input. However, the MFCC features in the LSTM model are only based on spectral information up to 4 kHz, whereas the DNN-HMM uses spectral representations of up to 8 kHz.
3. Facebook’s fairseq Wav2Vec 2.0 (Baevski et al., 2020) is a **CNN-Transformer** model that, in contrast to the previous two models, takes raw audio as input. Features are extracted using a convolutional neural network (CNN) that convolves the input signal over time. Temporal context is modelled using a Transformer architecture (Vaswani et al., 2017), which uses an attention mechanism to relate inputs from different timesteps.

We selected a battery of seven tests that measure psychometric curves under a range of distortions. These were chosen such that each test measures a conceptually different aspect of hearing (e.g. spectral content, temporal fine structure or speech in noise), and so that each relies on sentence- or word-based speech recognition. Phoneme and syllable recognition tasks were excluded because the ASR systems (which were trained to predict full sentences) performed poorly on such short nonsense segments. Note that our experiments were performed in a mismatched condition. That is, all models were trained using (mostly) clean speech, but were tested here on data that was manipulated in certain acoustic dimensions.

### 2.1 Spectral Invariance: Sensitivity to Bandpass Filtering

#### Background

Although humans can perceive frequencies from 20 Hz up to 20 kHz, even a few small frequency bands or a single moderately wide band can be sufficient to achieve high intelligibility in quiet. For example, Warren et al. (2000) showed that by using pass bands with a near vertical slope, high intelligibility rates are achieved when all but the 1 kHz to 2 kHz region is filtered out. Here, we investigate whether ASR systems can similarly make use of this ‘spectral redundancy’ in speech. As in Warren et al. (2000), speech signals were filtered with bandpass filters centered at 1500 Hz and a varying bandwidth (given in semitones, as in Warren et al. 2000).

#### Detailed comparison

Compared to human performance, the ASR systems required a much wider spectral range to perform well (Figure 1a). At the width at which humans have near-ceiling accuracy (12 semitones), the average accuracy was poor for all ASR systems (0-10% accuracy). Note that we used a different speech corpus for the ASR systems and humans in order to make our code freely accessible. The speech corpus we used to test the ASR systems is more difficult, but would only be expected to lead to a decrease of around 20% accuracy (see Signal Processing and Procedures for more details).

**Figure 1.**
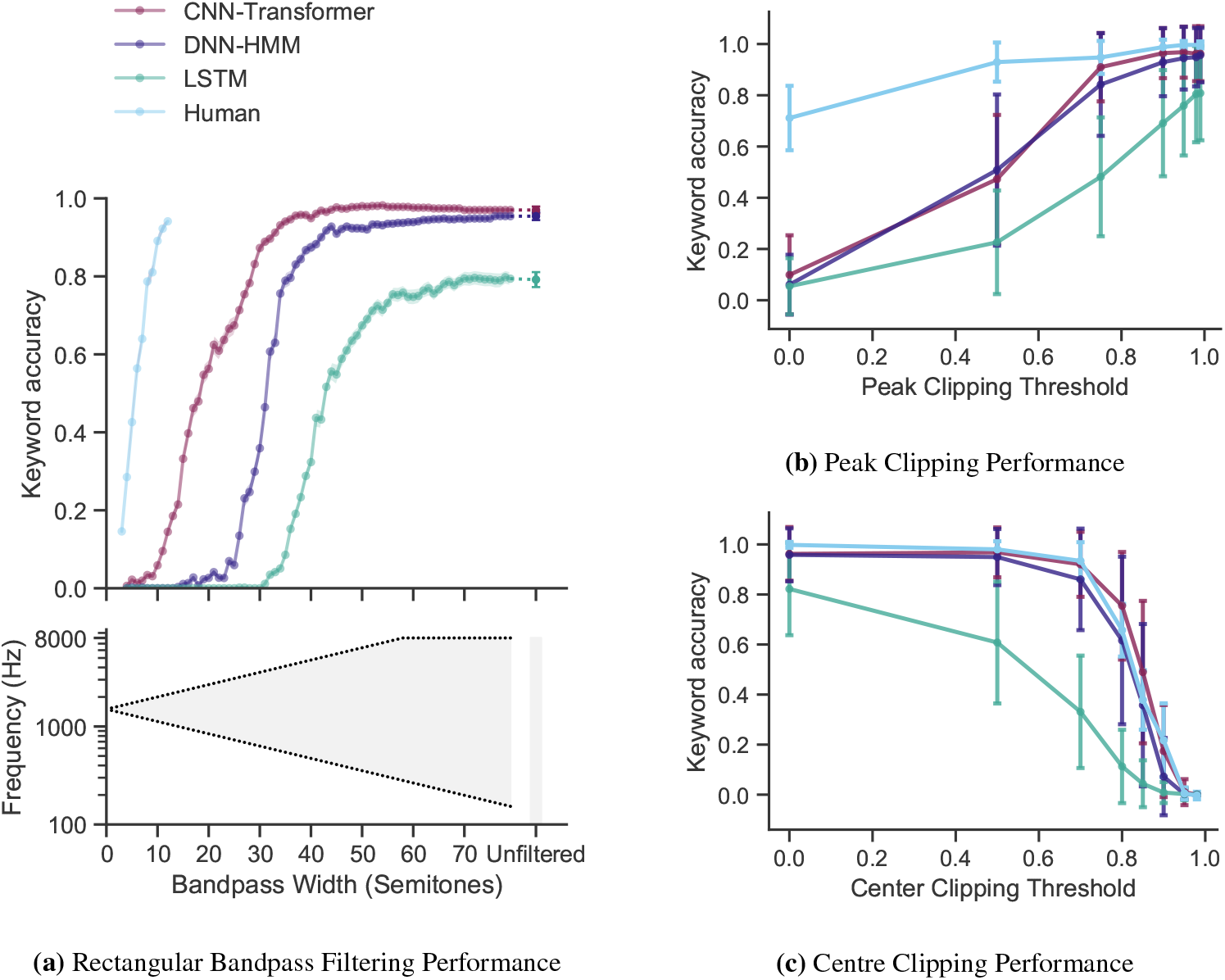
The performance of the three automatic speech recognition (ASR) systems compared to human performance for three types of distortions. Keyword accuracy denotes the percentage of correct keywords measured for 100 sentences. (**a**) Accuracy obtained when each sentence was bandpass filtered around 1500 Hz with a width that varied. Error bars denote the standard error of the mean. Below are the upper and lower cutoff frequencies of the filters. Human data from Warren et al. (2000). (**b**) Performance in peak clipping experiment. The peak clipping threshold *t* determines the percentage of the sentence samples that was not clipped. At *t* = 0 the sound is reduced to its signs. Error bars show one standard deviation. Human data from Kates and Arehart (2005). (**c**) As in b, but now using centre clipping, which means that all samples that were *lower* than the clipping value were set to zero. The centre clipping threshold determines the percentage of sentence samples that were *not* clipped. At *t* = 0 the original signal is retained. Error bars show one standard deviation. Human data from Kates and Arehart (2005).

There was a large variation between the ASR systems themselves. Despite the similar performance of the CNN-Transformer and DNN-HMM for unfiltered sounds, the CNN-Transformer was much more robust to bandpass filtering than the DNN-HMM. The LSTM, which had a lower accuracy in quiet than the two other models, was the least robust against bandpass filtering, with performance tapering off to near ceiling-level around 60 semitones compared to approximately 35 and 45 semitones for the CNN-Transformer and DNN-HMM, respectively. The MFCC input features of the LSTM model retain spectral information only up to 4 kHz (see Details of ASR Systems), which is reached by the bandpass filter at 34 semitones. At this level of filtering, performance was still close to zero for the LSTM, and performance did not increase drastically until around 40 semitones. This suggests that low frequency information between 40 and 60 semitones (265 Hz to 472 Hz) is particularly important for this model. The other two models, particularly the CNN-Transformer model, reached ceiling level well before such low frequency information was available.

### 2.2 Peak and Centre Clipping

#### Background

Communication systems such as microphones or loudspeakers can introduce a variety of nonlinear distortions. For example, when a microphone records very loud sounds it may saturate, resulting in the clipping of high amplitude pressure waves. This is referred to as peak clipping, a distortion that introduces additional higher harmonics as well as intermodulation products. Perhaps surprisingly, human speech recognition is robust against peak clipping distortions. Even when a speech signal is reduced to its signs (each sample replaced by +1 if positive or -1 if negative), normal-hearing humans achieve between 70% and 90% intelligibility (Licklider and Pollack, 1948; Kates and Arehart, 2005). A related distortion is centre clipping, which refers to the reduction of low-level amplitudes. This may occur when noise-suppression systems are in place. In humans, a small amount of centre clipping is not detrimental. However, higher levels of centre clipping affect intelligibility considerably, as weak consonants (which tend to have lower amplitudes than vowels) are stripped out (Licklider, 1946; Kates and Arehart, 2005). Following the approach described in Kates and Arehart (2005), we systematically introduced different levels of peak and centre clipping. This involved determining at which amplitude value *c* the absolute value of a fraction of *t* of the sound samples would be lower than *c*. We refer to *t* as the clipping threshold, a value that ranges between 0 and 1. For peak clipping, a sound is reduced to its signs for *t* = 0, whereas in the case of centre clipping the signal is reduced to only zeros at *t* = 1.

#### Detailed comparison

In both the peak and centre clipping experiments, the CNN-Transformer and DNN-HMM displayed similar robustness to one another, whereas the LSTM model performed much worse at all clipping thresholds (Figure 1b and 1c). This is in contrast to the bandpass filter experiment (Spectral Invariance: Sensitivity to Bandpass Filtering), where the CNN-Transformer outperformed the DNN-HMM. In the peak clipping experiment, none of the ASR models performed as well as humans (Figure 1b). At a peak clipping threshold of 0.5, the performance of humans was near ceiling, whereas the accuracy of the DNN-HMM and CNN-Transformer dropped to around 50%. At infinite peak clipping (*t* = 0.0), the performance of the latter two systems hovered around 10%, compared to 72% for humans. In the centre clipping experiment, both the CNN-Transformer and DNN-HMM intelligibility rates were similar to human intelligibility rates (Figure 1c), whereas the LSTM performance performed worse. This suggests that the CNN-Transformer and DNN-HMM, but not the LSTM, are robust against changes in the distribution of low-level amplitude parts of the speech signal.

Previous studies have reported similar detrimental effects of peak clipping on ASR performance. For example, Tachioka et al. (2014) reported 20% accuracy for a large vocabulary task (20% at *t* = 0.1), although the Gaussian Mixture Model (GMM) HMM ASR system in this study had low ceiling performance (60% at *t* = 0.9). Malek et al. (2016) reported an increase in Word Error Rate (WER) from around 10 % for undistorted speech to around 50 % for speech that was distorted through nonlinear amplification and clipping. This effect was observed across a range of different input features, including MFCCs, although ‘primitive’ Filter Bank Coefficients (FBCs) (i.e. Mel-scaled spectrograms with no cosine transforms) appeared slightly more robust.

Further research is required to understand why ASR systems perform poorly under peak-clipping distortions. On a rudimentary level, the poor performance may be influenced by the change in distribution of the acoustic waveform that peak clipping introduces. For the end-to-end CNN-Transformer, peak clipping affects the distribution of the input values the model observes, particularly after normalisation. For MFCCs (the input to the DNN-HMM and LSTM) peak clipping results in an increasing shift of the mean and reduction in covariance of the MFCC coefficients (Poorjam et al., 2017). Alternatively, poor performance on peak clipped audio may be a result of the specific way in which peak clipping distorts the spectral envelope, as it introduces extra energy at harmonics in the frequency spectrum. Since the ASR systems have a different response to changes in periodicity than humans do (see Target Periodicity), other changes in harmonic structure may affect them differently too. Lastly, it should be noted that peak clipping specifically retains certain temporal information (e.g. zero-crossings). The inability of the tested ASR systems to make use of temporal fine structure (see Temporal Fine Structure) may therefore also contributes to their poor performance on peak clipped audio.

### 2.3 Spectral and Temporal Modulations

#### Background

Speech is characterised by fluctuations of power in both time and frequency, which are referred to as temporal and spectral modulations. As shown in Spectral Invariance: Sensitivity to Bandpass Filtering, speech can remain intelligible even after drastic degradations in spectral content, depending on which frequencies are taken out. Conversely, it has been shown that even with very coarse temporal information, speech remains intelligible in the presence of sufficient spectral information (Drullman et al., 1994; Arai and Greenberg, 1998; Arai et al., 1999). To disentangle the importance of spectral versus temporal modulations, Elliott and Theunissen (2009) used a modulation filtering technique to remove specific spectral modulations (given in cycles/kHz) and temporal modulations (i.e. amplitude modulations, given in Hz). It was shown that in humans, most modulations can be removed without affecting intelligibility of speech in quiet. However, under noisy conditions temporal modulations of 1 Hz to 7 Hz and spectral modulations of 0 cycles/kHz to 1 cycle/kHz are necessary to obtain high intelligibility rates. In this section we explore to what extent ASR systems rely on modulations by following the notch filter experiment described in Elliott and Theunissen (2009). In this experiment, word accuracy was measured for sounds in which specific temporal and spectral modulations had been filtered out. Performance was measured in white noise with an SNR of 2 dB (as in Elliott and Theunissen (2009)) as well as at the lowest SNR at which near-ceiling performance was reached for the ASR systems without any modulation filtering (10 dB, 15 dB, 25 dB for the CNN-Transformer, DNN-HMM and LSTM, respectively, we refer to these as the high-SNR regime).

#### Detailed comparison

As with the bandpass experiment, the CNN-Transformer model was most robust against degradations, followed by the DNN-HMM model and the LSTM model (Figure 2). Although absolute performance levels varied depending on the ASR system and noise level, the relative importance of the spectral and temporal modulation regions was mostly consistent with the regions that are important for HSR. This was particularly evident when the SNR levels were adjusted for near-ceiling performance level (the high-SNR regime, dotted lines in Figure 2).

**Figure 2.**
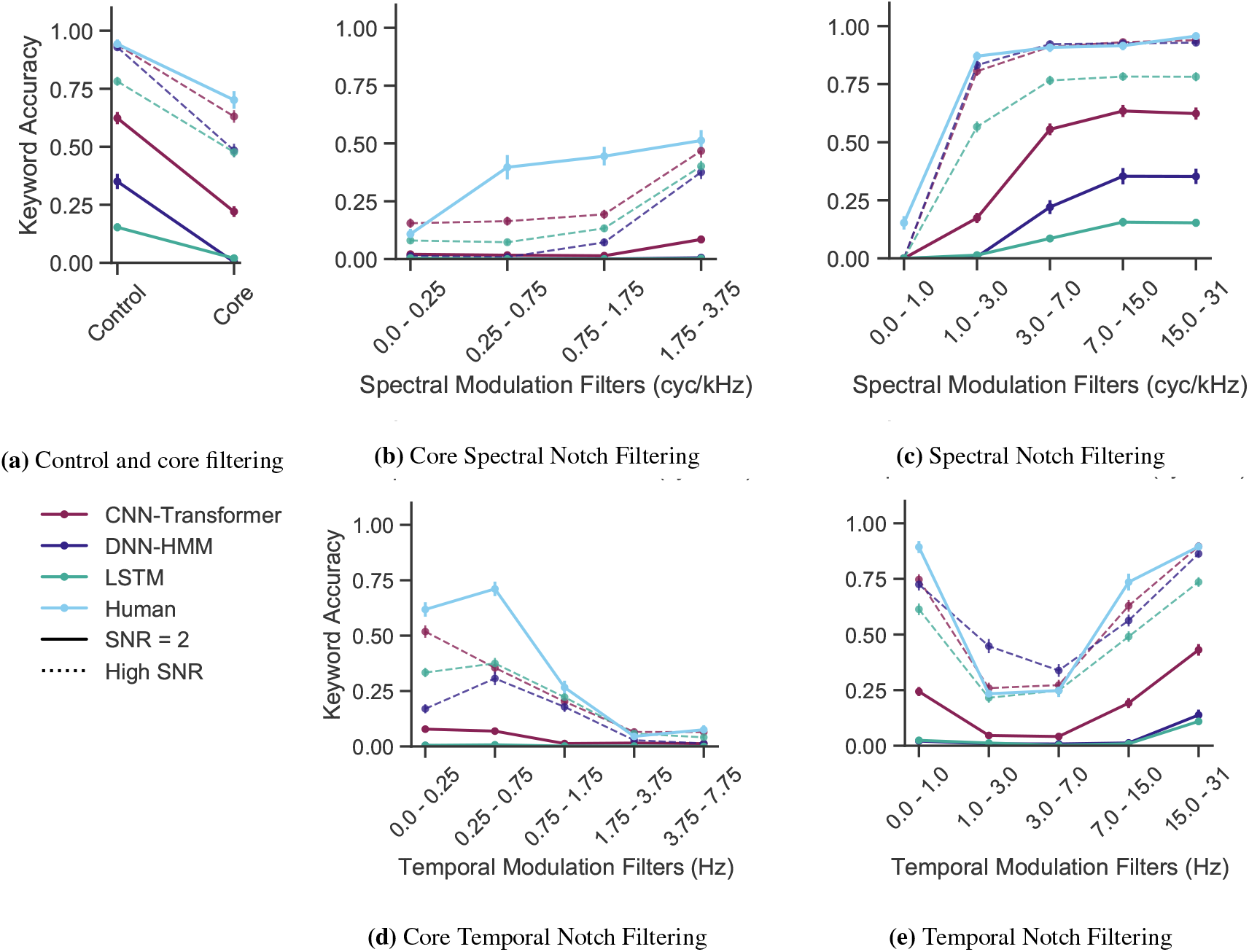
The performance of the three ASR systems in the spectral and temporal modulation filter experiment. Either spectral or temporal modulations are removed by computing the modulation power and phase spectra, setting specific regions to 0 and inverting. Performance is measured as percentage of keywords correct in 100 sentences embedded in white noise with a low (solid lines) or high (dashed lines) signal to noise ratio. Note that the maximum performance in quiet of the LSTM was only 80%. Error bars are plotted as the standard error of the mean. Human reference from Elliott and Theunissen (2009). (**a**) Performance in the control condition (spectrogram inversion without modulation filtering) and core region condition (removal of all spectral modulations above 3.75 cycles/kHz and temporal modulations above 7.75 Hz). (**b d**) Performance when, in addition to removal of all modulations outside of the core region, either spectral modulations (b) or temporal modulations (d) in the given regions are removed. (**c, e**) Performance when spectral (c) or temporal (e) modulations in the given regions are removed.

Similar to humans, ASR performance was most disrupted by the removal of low spectral modulations in the 0-1 cycles/kHz region, after which performance monotonically increases as the notch region covers higher spectral modulations (Figure 2c). In humans, performance in all other regions is near ceiling, but in the ASR systems the 1-3 cycles/kHz region is more important than the higher regions. The 0-1 cycles/kHz and 1-3 cycles/kHz regions roughly correspond to vowel formant separation and formant transitions in English (Elliott and Theunissen, 2009).

In both the ASR systems and humans, temporal notches in the 1-7 Hz region degrade intelligibility most, followed by notches in the 7-15 Hz region (Figure 2e). The regions roughly correspond to syllable and phoneme rates, which respectively are around 2 Hz to 5 Hz (Pickett, 1999) and 15 Hz to 30 Hz (Liberman, 1970; Greenberg et al., 1999). In humans, the removal of both very slow (0-1 Hz) and fast (15-31 Hz) temporal modulations does not lead to a significant reduction of intelligibility compared to unfiltered speech (control). For all ASR systems, on the other hand, removing the 0-1 Hz range does lead to a reduction in performance. Filtering these slow temporal modulations leads to fluctuating sound levels across the sentence with a rate of up to 1 Hz. Our results thus suggest that the ASR systems, unlike humans, are not able to adaptively respond to these slow changes in sound level.

When all modulations outside of the ‘core’ modulations (defined as up to 3.75 cycles/kHz and 7.75 Hz) are removed, humans retain more than 70% accuracy at a 2 dB SNR (Figure 2a). At the same SNR, ASR performance is much lower. In the higher SNR regime, the core region performance of the ASR systems is more comparable to that of humans. The results based on additional filtering in the core region provide a more refined view of the differences in sensitivity to slow spectral and temporal modulations. This shows that the ASR systems are more sensitive to spectral modulations in the 0.25-1.75 cycles/kHz region (Figure 2b). Furthermore, in humans the effects of temporal modulations in the 0.25-0.75 Hz region are not significantly different from those in the 0.0-0.25 Hz region (Figure 2d). For the CNN-Transformer, there is a relatively higher sensitivity to the 0.0-0.25 Hz range, whereas the shape of the DNN-HMM and LSTM are more comparable to that of humans.

In summary, the three ASR systems appear to be less robust against the white noise masker than humans are, but display roughly the same relative importance of spectral and temporal modulations. The similarity between human and ASR results contribute to the universality of the ‘speech modulation transfer function’ proposed in Elliott and Theunissen (2009). Furthermore, despite the use of a linear frequency scale in the computation of the MPS in Elliott and Theunissen (2009) (which differs from the approximate log scale at which both the human auditory periphery and MFCC feature extraction operate), these results further emphasise the importance of the use of ASR input features that retain (whether explicitly or implicitly) critical temporal and spectral modulations.

### 2.4 Temporal Fine Structure

#### Background

In the human auditory system, incoming sounds are decomposed into a set of narrow frequency bands by the cochlea. The responses in each frequency channel can be described through two components that function at different time scales. The slower component, the temporal envelope, reflects slow fluctuations in amplitude, whereas the rapid fluctuations of the narrowband carrier are referred to as temporal fine structure (TFS). In quiet listening conditions, speech with disrupted TFS that retains channel envelope information is highly intelligible. However, TFS is thought to play a more important role in challenging listening environments. For example, Hopkins and Moore (2010) showed that for normal-hearing listeners TFS information contributes significantly to the perception of speech in the presence of a competing talker, particularly for channels with centre frequencies below 1 kHz. Such a benefit was not observed for hearing-impaired listeners. As MFCC features do not retain detailed timing structure, the use of TFS information in the DNN-HMM and LSTM is likely very limited. The CNN-Transformer model, on the other hand, would at least in principle be able to leverage TFS information. However, considering that the CNN-Transformer model was not trained in noise, it may still have learned to rely solely on envelope cues. To test these hypotheses, we follow the two experiments described in Hopkins and Moore (2010). The first experiment explores the impact of cumulatively adding intact TFS starting from high or low frequencies (referred to as the *TFS-low* and *TFS-high* conditions, respectively). The second experment measures the impact of removing envelope and TFS information in specific spectral regions.

#### Detailed comparison

In humans, an increase in TFS information (in both the *TFS-low* and *TFS-high* conditions) leads to an increase in performance (reduction of SRT), particularly when TFS information is added to the low-frequency channels below 1 kHz (Figure 3). The overall SRT improvement from speech that contains mostly envelope information to unproccesed channels is around 6.6 dB. The absolute SRTs observed in the ASR systems are much higher, which can partly be explained by the use of a different dataset and partly due to reduced robustness of ASR systems in noise. For all ASR systems, there is some improvement with the addition of intact TFS information, but the overall benefit is much smaller (around 0.5 dB, 1 dB, 2 dB for the CNN-Transformer, DNN-HMM and LSTM, respectively) and there does not appear to be a particular preference for low-frequency TFS regions. Disrupting the TFS information also somewhat changes the envelope, so the observed small benefit of intact TFS is likely to be a consequence of better preservation of the envelope.

**Figure 3.**
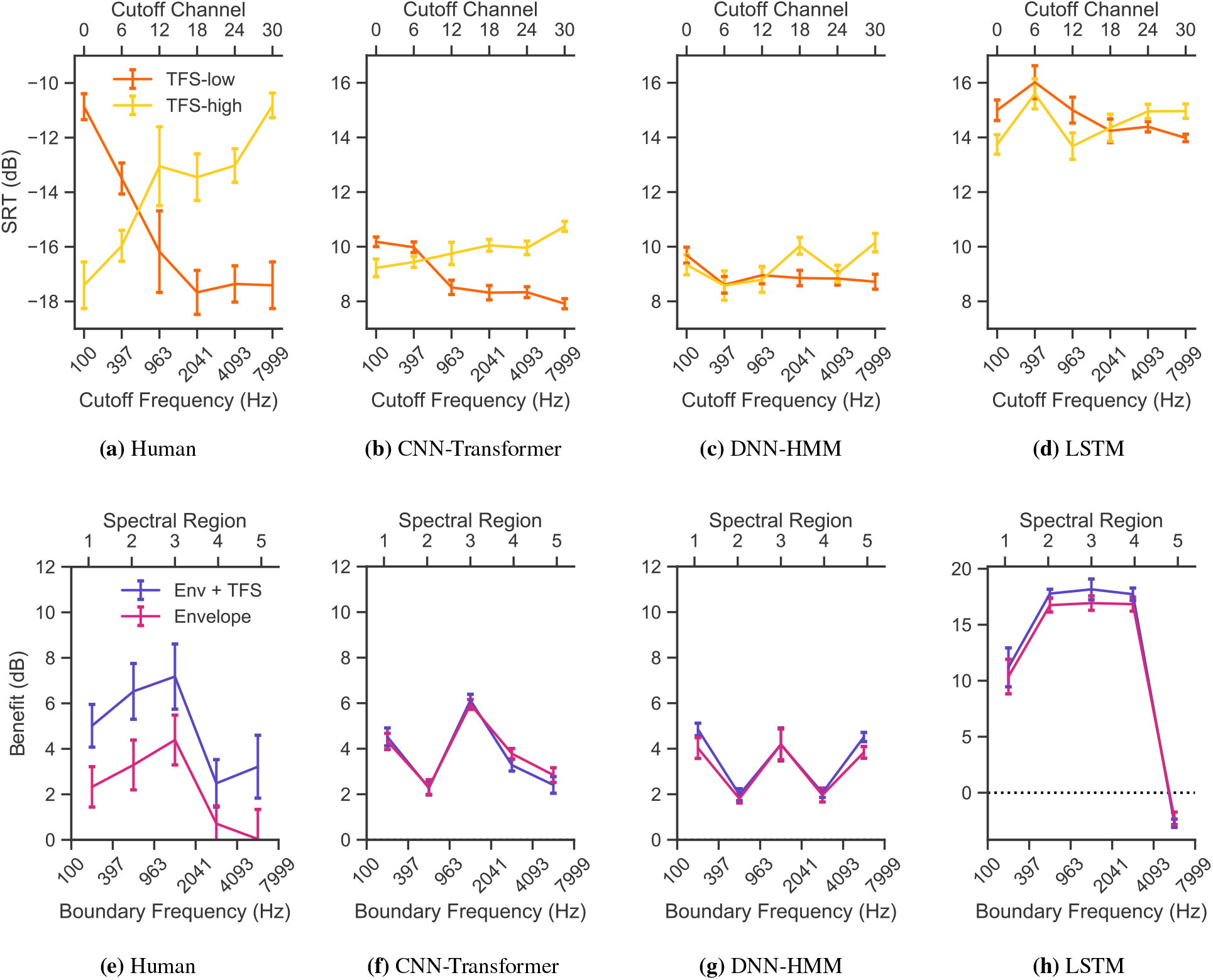
Performance in the two temporal fine structure (TFS) experiments measured for 100 IEEE sentences (20 per SRT, 5 per condition) in six-speaker babble noise. In these experiments, TFS is either distorted or retained in specific frequency channels using a tone vocoder. This vocoder initially filters speech in 30 frequency channels. To distort TFS from a given channel, the channel response is replaced by a tone that is modulated by the channel envelope. Human reference data, which was measured on the CRM task corpus in single-speaker babble noise, were taken from Hopkins and Moore (2010). Error bars are plotted at standard error of the mean. (**a-d**) In the first experiment, intact TFS information is added cumulatively starting either from the highest (*TFS-high*) or lowest (*TFS-low*) spectral region. The SRT-70s are plotted as a function of cut-off region for each of these conditions. Note that at both the highest cut-off frequency in *TFS-high* and lowest cut-off frequency in *TFS-low*, the whole sentence is vocoded. Conversely, TFS information across the whole frequency spectrum is present at the lowest *TFS-high* region and highest *TFS-low* region. (**e-h**) In the second experiment, TFS information was either distorted or retained in a given spectral region, while all other regions were vocoded. A third condition was also tested in which the spectral region was completely removed. The benefit of adding envelope information was calculated as the difference between the SRT for the condition in which all regions were vocoded (i.e. TFS was removed in that region) and the SRT for the condition in which the spectral region was removed from a fully vocoded sentence. The benefit from adding envelope and undistorted TFS information together was calculated as the difference between SRT for the condition in which the spectral region was removed fully and the condition in which TFS was retained.

In the second experiment, a comparison is made between the benefit of adding both envelope and intact TFS information or envelope information alone to a specific spectral region. The SRT benefit is determined by taking the difference in SRT between a vocoded sentence in which a specific spectral region is completely removed and the SRT of a sentence in which the envelope of that region is retained (envelope benefit) or in which both the envelope and TFS are retained (envelope + TFS benefit). In normal-hearing listeners, the addition of TFS information leads to a benefit between 2 dB to 3 dB across all spectral regions. In contrast, the addition of intact TFS information does not lead to an improvement in performance for any of the ASR systems. This is consistent with the results of the first experiment, where no large difference was observed when incorporating intact TFS information from a single region. The results indicate that the spectral regions are too fine-grained to obtain an observable benefit in ASR systems. The second experiment also exposes the differences in the relative importance of the spectral regions, in line with the results found in Spectral Invariance: Sensitivity to Bandpass Filtering. In humans the removal of envelopes in the high frequency regions (above 2 kHz) does not affect performance much. The ASR systems, on the other hand, display significant drops in performance when these regions are removed. Compared to the CNN-Transformer and the DNN-HMM, the LSTM model relies more heavily on the second, lower-frequency region (400 Hz to 1000 Hz), something that was also observed in Spectral Invariance: Sensitivity to Bandpass Filtering. Note that the LSTM does not benefit from the spectral region 4 kHz to 8 kHz since the input MFCCs to this model are based on spectral information up to 4 kHz only.

In summary, in contrast to normal hearing humans, the ASR systems do not use TFS information and rely more heavily on the presence of envelope information across the entire frequency spectrum than humans do.

### 2.5 Target Periodicity

#### Background

A speech sound generally consists of both periodic and aperiodic segments. Periodicity in speech is caused by periodic vibrations of the vocal cords, as can be heard in voiced speech. Aperiodicity in speech results from constrictions in the vocal tract, for example in fricatives. Periodicity, which is closely associated with pitch, likely plays an important role in extracting certain linguistic features such as intonation. In a systematic comparison between vocoded speech with artificially varied levels of periodicity, it was shown that in quiet listening conditions the presence of aperiodic segments is important for speech intelligibility (Steinmetzger and Rosen, 2015). In particular, noise vocoded speech (which has no periodicity) is about as intelligible as speech with a natural balance of periodicity and aperiodicity (also called Dudley-vocoded speech), whereas fully periodic speech tends to be less intelligible. This effect is particularly noticeable when the spectral resolution is reduced due to a smaller number of vocoder channels. To further understand the use of periodicity in speech in ASR systems, we tested the three ASR systems on the same sentences that were used in the experiment of Steinmetzger and Rosen (2015).

#### Detailed comparison

Steinmetzger and Rosen (2015) measured accuracy for three types of vocoded speech with up to 16 channels in quiet. It was shown that for humans, the absence of any periodicity information (and hence intonation) does not significantly affect speech intelligibility in quiet listening environments (Figure 4). On the other hand, additional unnatural periodicity cues as found in the fully periodic vocoded speech leads to substantially poorer speech intelligibility. In Figure 4, we also included estimates of performance in quiet based on psychometric functions for 24 channels as well as TANDEM-STRAIGHT (TS) speech from a subsequent experiment (see Masker Modulations and Periodicity). TANDEM-STRAIGHT is a general-purpose speech analysis and synthesis system that can be used to manipulate aspects of periodicity in speech without reducing its spectral resolution. The differences between the three levels of periodicity mostly disappear when using the more natural sounding TANDEM-STRAIGHT speech sounds, which were produced to be either not periodic (noise-vocoded), fully periodic or mixed-periodic.

**Figure 4.**
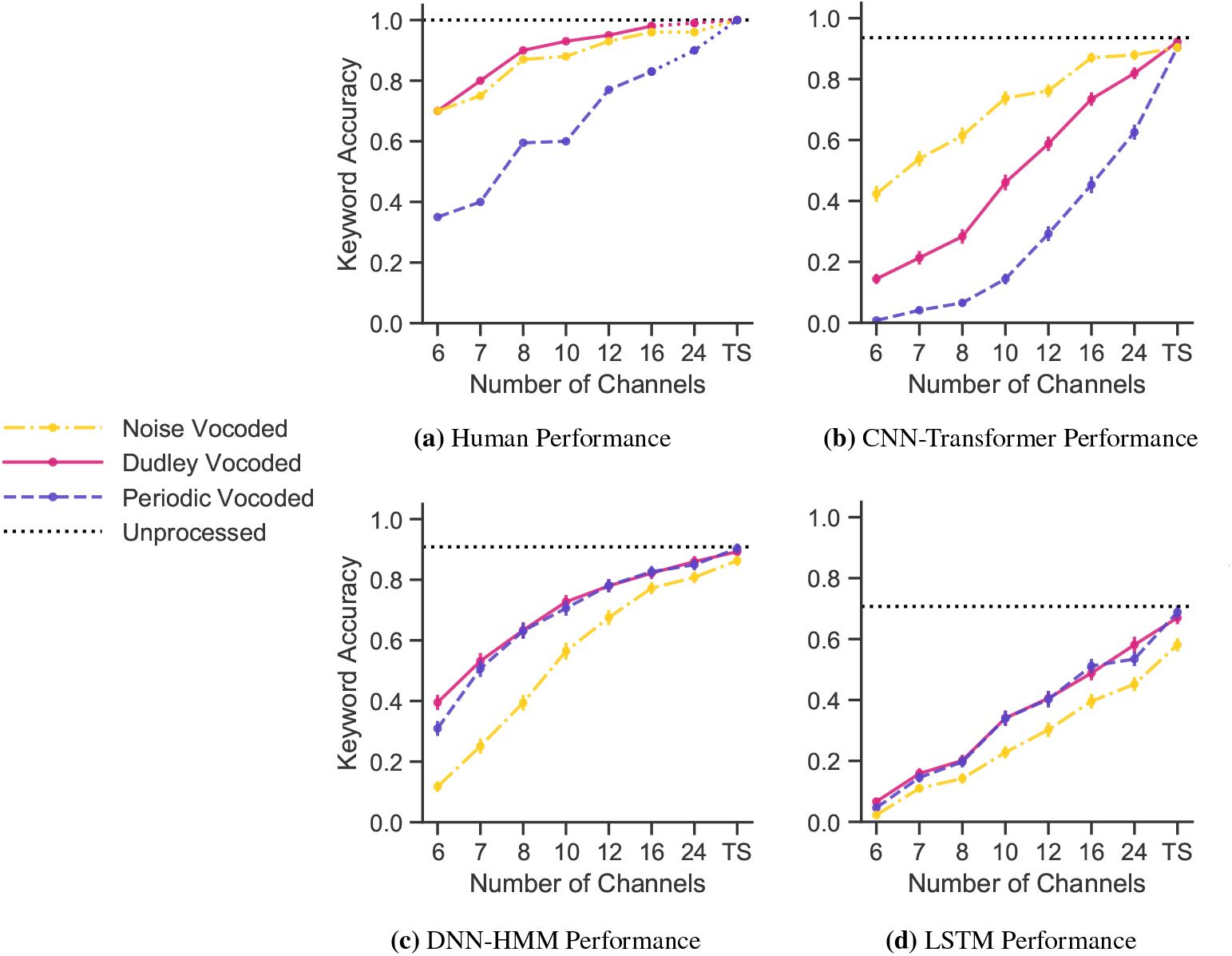
Performance in the target periodicity experiment. In this experiment, recognition accuracy is measured in three vocoding conditions that differ in periodicity: noise vocoding (aperiodic), Dudley vocoding (mixed-periodic) and fully periodic. The TANDEM-STRAIGHT (TS) condition, in which the periodicity of the source was controlled using the TANDEM-STRAIGHT algorithm, is also included. Performance is measured as the percentage of correct keywords and is plotted as a function of the number of frequency bands for the three different vocoding conditions. The accuracy for unprocessed speech is denoted by the dashed horizontal line. All error bars show the standard error of the mean. (**a**) Human performance taken from Steinmetzger and Rosen (2015). Performance in the 24-channel vocoded and TS conditions was estimated from psychometric function curves from a second experiment that investigated the effects of noise on speech with varying levels of periodicity. (**b-d**) Performance of CNN-Transformer (**b**), DNN-HMM (**c**) and LSTM (**d**) on 100 IEEE sentences presented at 65 dB. The material was identical to the material used in Steinmetzger and Rosen (2015).

None of the ASR models show the same relative sensitivity to periodicity in speech. For example, in stark contrast with the poor intelligibility for periodic speech found in humans, the DNN-HMM and LSTM models perform equally well with respect to Dudley and fully periodic vocoded speech. For both these models, noise vocoded speech leads to the poorest performance and it remains lower than for the other two speech types even for the more natural sounding TANDEM-STRAIGHT speech. Since these two models both use MFCCs as input, these may be the source of the discrepancy. Although MFCCs discard most periodicity information, some voicing information is retained (Milner and Shao, 2006). Our results suggest that both the DNN-HMM and LSTM model rely on the presence of these voicing cues (which are mostly retained in the higher MFCC coefficients) in making predictions. In contrast to the DNN-HMM and LSTM, the CNN-Transformer does display a decrease in performance for periodic speech compared to dudley-vocoded speech, which suggests that the lack of periodicity is used as a cue. Surprisingly, however, the CNN-Transformer performs best when noise-vocoded speech is used. It may be the case that certain mismatches between periodic and non-periodic components that result from the imperfect automatic *F*_0_ estimation of Dudley-vocoded speech negatively affect performance. As in humans, performance for the TANDEM-STRAIGHT speech converges to that for unfiltered speech.

### 2.6 Competing Talker Backgrounds

#### Background

One of the most challenging listening environments is a background of other speakers. Babble maskers affect speech intelligibility in at least two ways. Firstly, ‘energetic masking’ (EM) may occur, caused by the presence of masker energy in the same frequency region(s) as the energy in the target signal. As speech is a modulated signal, the amount of EM will vary over time, with short glimpses during which the SNR is much higher. These glimpses may either be comodulated, as occurs with modulations consistent across the spectrum, or uncomodulated, meaning they are restricted in frequency. There is evidence that some ASR systems take advantage of at least comodulated dips to process speech in noise (Spille and Meyer, 2017). A second type of masking is ‘informational masking’ (IM), which occurs when the target speech and masker are difficult to disentangle despite a lack of energetic overlap, for example because the masker has a similar voice quality, or the linguistic content of the masker is distracting (lexical interference). To investigate the relative importance of EM and IM, Rosen et al. (2013) used babble noise, noise-vocoded and envelope-modulated speech-shaped noise with an increasing number of competing talkers. Each of these maskers may introduce varying levels of EM and IM. For example, the masker effectiveness of a babble masker initially increases in going from one competing talker to two competing talkers, but slowly decreases after more than two speakers are added. The latter decrease suggests a reduction in IM, resulting from the individual talkers ‘blending’ together and therefore reducing intelligibility of individual speakers. This release is large enough to counteract the reduced glimpsing opportunities (i.e. increase of EM) that occur when the number of talkers increases. A similar influence of IM was not present for the noise-vocoded speech, which may be a result of the differences in sound quality and periodicity. The modulated noise masker, which can induce comodulated EM, is the least effective masker when it is modulated at the rate of a single speaker. However, masker effectiveness improves as the number of speakers increases. To investigate the way background speakers affect ASR performance, we tested the ASR systems following the experimental paradigm of Rosen et al. (2013).

#### Detailed comparison

Figure 5 shows the SRT-50s obtained for speech in the three maskers (babble, noise vocoded babble and noise modulated with the envelope of the babble). Although the SRT-50s observed for the ASR systems were much higher than those for humans, two of the three main results presented in Rosen et al. (2013) are reflected in the results of the ASR systems, particularly the DNN-HMM.

**Figure 5.**
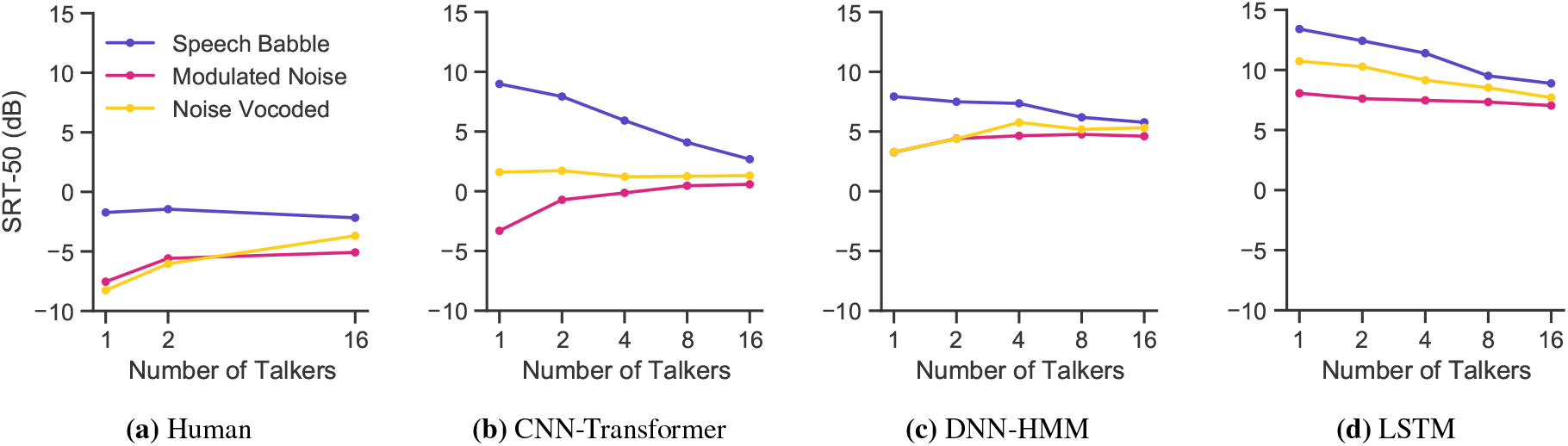
Performance in the competing background talkers experiment. Speech reception thresholds (SRTs) at 50% correct for 100 IEEE sentences were measured in three different types of maskers: speech babble, noise-vocoded babble and modulated speech-shaped noise. These maskers were created with a varying number of talkers (1, 2, 4, 8 and 16). SRTs are plotted as a function of the number of talkers in the masker. (**a**) Human data taken from Rosen et al. (2013). (**b-d**) Performance of the CNN-Transformer (**b**), DNN-HMM (**c**) and LSTM (**d**) ASR systems on the same material as used in Rosen et al. (2013).

Firstly, for all ASR systems babble was always the most effective masker, as it had the highest SRT-50. For humans, SRTs using noise-vocoded babble and modulated noise maskers tend to closely follow one another, a result that was only clearly replicated by the DNN-HMM.

Secondly, for humans performance for the modulated and noise-vocoded maskers tends to increase monotonically, with an initial rapid increase after which the SRT-50 plateaus. Although less pronounced, a similar effect was observed for the DNN-HMM for both types of noise. For the CNN-Transformer, this only occurred for modulated noise, as the SRT-50s for noise-vocoded speech remain constant. As shown in Target Periodicity, the CNN-Transformer had high intelligibility rates for noise-vocoded speech. Together these results suggest that the potential benefit of a reduction of EM is counteracted by informational masking. In contrast to the other two ASR systems, the SRT-50s using modulated noise did not change as the number of speakers increases for the LSTM. This suggests that this model is not able to make use of glimpsing opportunities.

Lastly, a major difference between the ASR systems and the human outputs are the single babble speaker effects. Rosen et al. (2013) report that the SRT-50 for maskers with one and two speakers is almost identical, but report that single-speaker masker was a less effective masker at lower SNRs than two-speaker babble. Such a strong single speaker benefit was not observed for the ASR systems. One explanation for this is that the humans were instructed to focus on one specific target speaker. The ASR systems, on the other hand, will likely transcribe parts of both sentences or even only the masker speaker at low SNRs.

In summary, the DNN-HMM and CNN-Transformer, but not the LSTM model, displayed similar interactions of IM and EM as humans do. However, in contrast to humans, performance for single-speaker maskers was much worse. Furthermore, the CNN-Transformer showed a reduced ability to make use of glimpsing opportunities when exposed to the noise vocoded masker, likely as a result of IM introduced by the noise vocoder, which (as was shown in Target Periodicity) was highly intelligible to the CNN-Transformer.

### 2.7 Masker Modulations and Periodicity

#### Background

In Competing Talker Backgrounds, our discussion mostly focused on the effects of masker modulations and masker intelligiblity on speech in noise perception. However, another major difference between a speech and noise-vocoded speech masker is the level of periodicity, which is present in a speech masker but absent in noise vocoded speech. In Target Periodicity, it was shown that ASR systems respond differently to periodicity in the target speaker. In this section, we compare the effects of periodicity in a masker with masker modulations following the approach described in Steinmetzger and Rosen (2015). They compared the advantage of using a periodic masker (masker periodicity benefit; MPB) with the advantage from modulations in the masker (fluctuating masker benefit; FMB) and found that the MPB tended to be higher. As in Steinmetzger and Rosen (2015), we compare two types of noise (speech-shaped noise and harmonic complexes) that are either modulated or unmodulated. The SRT for speech in each of these four maskers is determined for the same three types of speech (noise-vocoded, mixed-periodic and fully periodic) as described in Target Periodicity.

#### Detailed comparison

The amount of MPB and FMB in humans depends mostly on the intelligibility rather than the type of target speech. For example, the MPB for near-ceiling performance is around 8 dB regardless of the type of vocoding (Figure 6). Similarly, the FMB hovers between 2 dB to 5 dB, with the benefit being slightly higher for noise maskers than for harmonic complex maskers. The performance of the ASR systems similarly did not show a clear effect of periodicity in the target speech, as exemplified by the MPBs and FMBs for the TANDEM-STRAIGHT sentences (Figure 6). However, only the CNN-Transformer model displayed a clear MPB and FMB. The LSTM and DNN-HMM, on the other hand, had a negligible MPB and a negative FMB, suggesting that masker modulations interfere with speech in noise performance.

**Figure 6.**
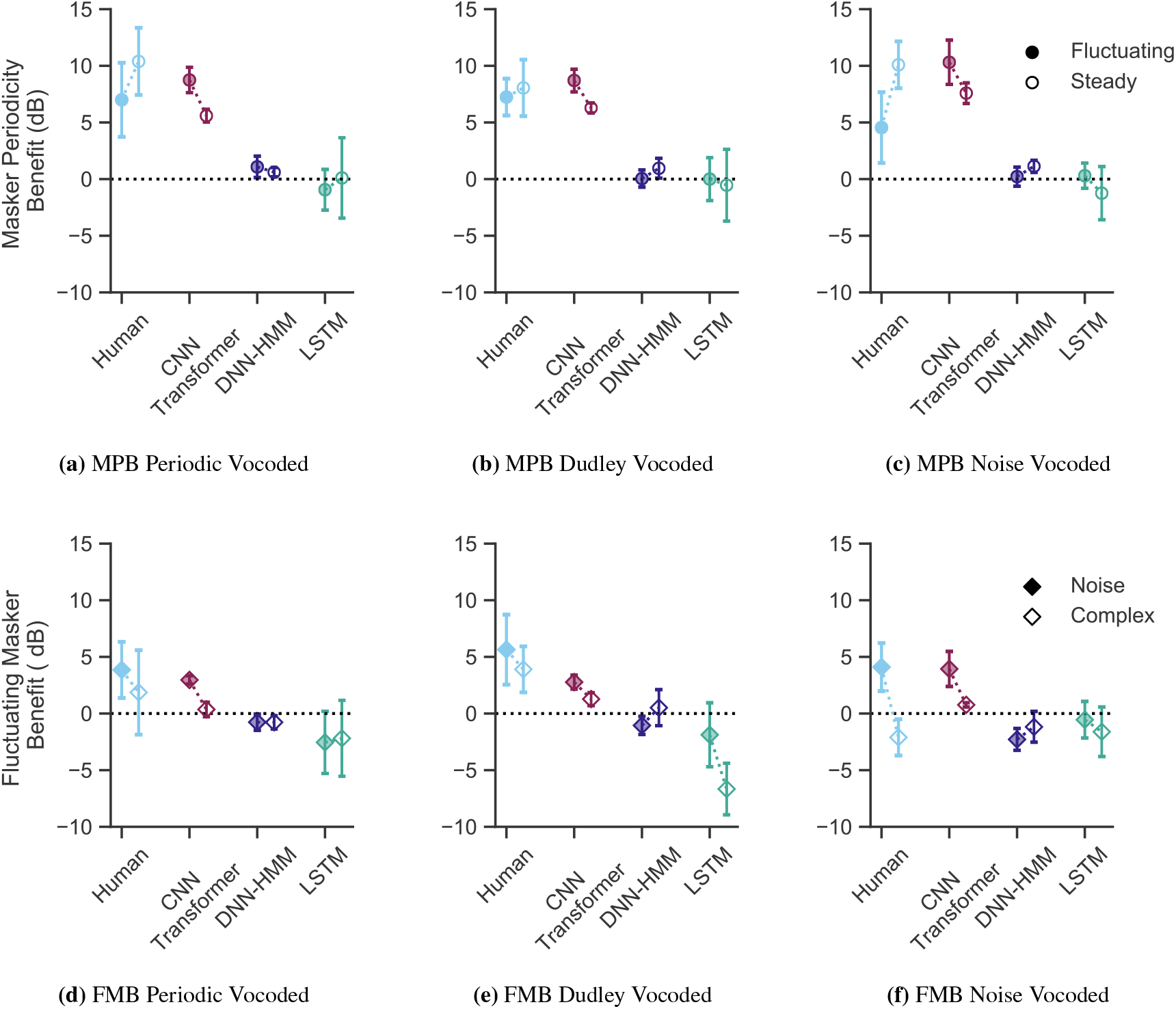
The obtained median masker periodicty benefits (MPBs) and fluctuating masker benefits (FMBs) in the masker periodicity experiment. In this experiment, SRTs were measured for speech in both noise and harmonic complex maskers that were either modulated or not. The target speech had different levels of periodicity (see Figure 4), namely fully periodic (**a**,**d**), mixed periodic (Dudley vocoded, **b**,**e**) and non-periodic (noise vocoded, **c**,**f**). These were created using TANDEM-STRAIGHT, a procedure that results in natural sounding stimuli. Human data were taken from Steinmetzger and Rosen (2015), and the ASR system performance was measured on the same stimuli as for the human data (100 IEEE sentences, measured in blocks of 20 sentences). All SRT-50s were determined using a static SRT method (see Speech Reception Threshold Determination). Error bars denote standard deviation. (**a-c**) MPBs are computed as the difference between the SRT-50s obtained with steady and modulated maskers. (**d-f**) FMBs denotes the difference in SRTs between the and periodic (noise) and non-periodic (harmonic) maskers.

For humans, higher intelligibility is associated with both higher MPB and higher FMB (Figure 7). The benefit for periodic maskers is always positive, but modulated maskers interfere if the target speech intelligibility is low. Of the ASR systems, only the CNN-Transformer showed both trends. The DNN-HMM model gave a very small MPB of around 1 dB, but the benefit did not increase with an increase in target speech intelligibility. Both the LSTM and DNN-HMM models showed an upward trend in FMB, suggesting that the modulated masker provides less interference when intelligibility is higher. However, neither achieved a positive benefit.

**Figure 7.**
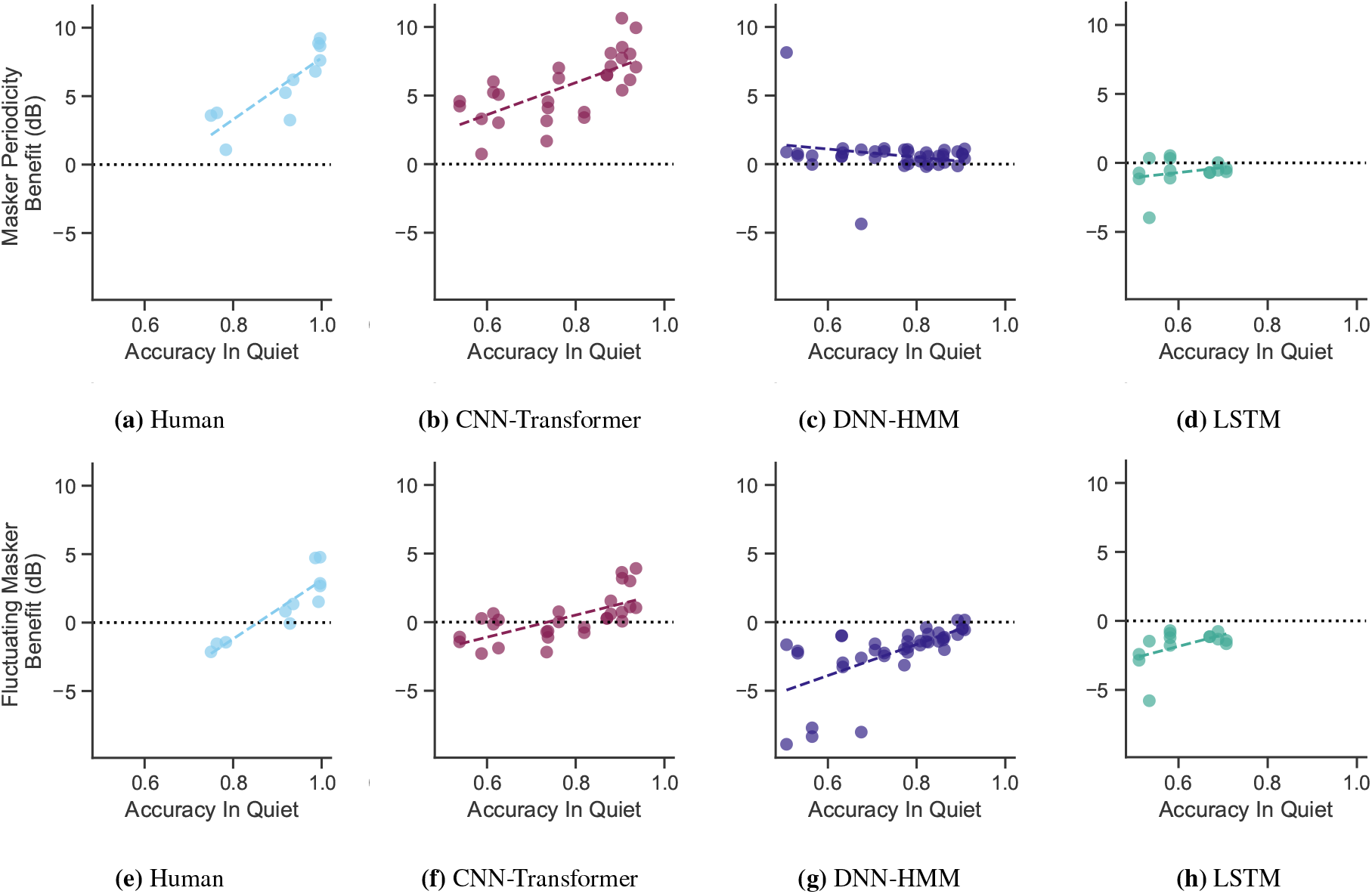
Results from the masker modulations and periodicity experiments measured over a range of target speech. Here, the masker periodicity benefit (MPB) (**a-d**) and fluctuating masker benefit (FMB) (**e-h**) are plotted as a function of performance for the target speech in quiet for humans (**a, e**), the CNN-Transformer (**b, f**), the DNN-HMM (**c, g**) and the LSTM (**d, h**). Accuracy in quiet was measured as in the target periodicity experiment (see Figure 4). Datapoints reflect performance for the three target vocoding types (periodic, Dudley vocoded and noise vocoded) with a varying number of vocoded channels, the TANDEM-STRAIGHT (TS) vocoded speech as well as unprocessed speech. The MPB and FMB were computed as described in Figure 6, and are based on SRT-50s pooled over all 100 IEEE sentences. Human data are based on the median values presented in Steinmetzger and Rosen (2015). For the TS and unprocessed speech, performance in quiet was estimated from the psychometric curves presented in that work.

Modulations in the masker thus only appears to provide a benefit for the CNN-Transformer, in line with that observed in the speech in babble experiment (see Competing Talker Backgrounds). The lack of modulation benefit for the DNN-HMM is somewhat surprising, since the results of the speech in babble experiment suggest a small benefit of around 1 dB for noise that is modulated by the envelope of a single-speaker compared to noise modulated by 16-speaker babble. It may be the case that the DNN-HMM requires larger gaps than those introduced by the 10 Hz amplitude modulations used in this experiment. Since the average syllable rate in English ranges from 2 Hz to 5 Hz, speech-modulated noise may provide more glimpsing opportunities. Previous work has shown that a similar DNN-HMM model was able to make use of glimpsing opportunities in noise maskers that were modulated with a rate of 8 Hz, although that model was trained in modulated noise (Spille and Meyer, 2017).

Steinmetzger and Rosen (2015) proposed several factors that could explain the presence of the periodicity benefit for humans. Firstly, harmonic complexes may allow for ‘spectral glimpsing’ in between the individual harmonics of the complex maskers. As shown in Competing Talker Backgrounds, the DNN-HMM and the CNN-Transformer showed similar patterns of sensitivity to glimpsing opportunities in noise-vocoded and modulated noise. This may account for the small periodicity benefit observed for the DNN-HMM. Secondly, periodic maskers typically hardly fluctuate outside of the *F*_0_ modulation range itself, particularly not at the low modulation rates that are most important for speech intelligibility. However, if this were the most important factor, we would expect both the DNN-HMM and LSTM model to benefit from this too, since these two models are most sensitive to the presence of slow temporal modulations (see Spectral and Temporal Modulations). Lastly, harmonicity in the masker could enable the auditory system to effectively subtract the masking sound from the signal mixture (de Cheveigné et al., 1995). For humans, the mixed target provides less MPB than the fully periodic or noise-vocoded targets. It may be that the interruptions in periodicity in the mixed target make it more difficult to form two separate auditory streams, whereas it is easier to cancel out the harmonic background in the presence of a fully aperiodic target sound or to follow two separate periodicities. Since the DNN-HMM and LSTM models are more sensitive to the presence of periodicity at lower spectral resolutions, it may be the case that the addition of another periodic signal is more disruptive. In contrast, the CNN-Transformer model is best at recognising noise-vocoded speech at lower spectral resolutions. The focus on unvoiced speech segments may allow this model to cancel out the harmonic complexes more easily or ignore them entirely.

### 2.8 Model Retraining

We tested “off the shelf” ASR systems that were not trained with examples of the distorted test speech. In common with most machine learning systems, ASR systems rely on the assumption that the training and evaluation data come from the same distribution. It is therefore unclear whether a reduction in performance for distorted speech is the result of the distribution shift or because the model truly relied on the feature that was distorted. By contrast, in most of the experiments described, the human listeners were given time to familiarise themselves with the distorted speech material. Unfortunately, it is not possible to create a perfectly fair equivalent, since most machine learning systems require much more data than humans in order to learn. This makes it difficult to “familiarise” an ASR system without extensive retraining far beyond that of a human listener. To somewhat mitigate this disadvantage, we avoided distortions that humans would have been trained to listen to from a young age, such as whispered speech. However, in some of our experiments humans still have a clear advantage. For example, comparing these ASR systems with speech in competing speech is something humans have been exposed to many times before. In many ways, it is surprising that some of the ASR systems perform so similarly to humans at specific tasks without being trained to do so.

It would be tempting to think that retraining the models with examples of distorted speech would lead to closer matches. However, we found that retraining to improve performance in one task can lead to worse performance at a different task. To illustrate this, we retrained smaller versions (i.e. with fewer parameters) of the CNN-Transformer model with either a dataset of only unfiltered speech, or a dataset in which 50% of the data was bandpass filtered with randomly selected bandwidths. We chose to finetune on the ‘BASE’ CNN-Transformer, a version of the model with only 12 rather than 24 Transformer blocks, due to resource constraints. Our finetuning procedure followed the finetuning parameters and approach described in Baevski et al. (2020) for finetuning the ‘BASE’ model on 100 h of the LibriSpeech dataset. As expected, the models trained on bandpass filtered data performed better in the bandpass filter test than the models trained with unfiltered speech data (Figure 8a), even outperforming the bandpass filter performance of the much larger CNN-Transformer model that was trained on unfiltered speech (Figure 1a). The model accuracy is comparable to that of humans tasked with recognising bandpass filtered isolated words (Stickney and Assmann, 2001). However, further testing showed that while spectral invariance improved, noise robustness in the retrained model was decreased (Figure 8b and 8c). In particular, the models trained on bandpass filtered speech have consistently higher SRT-50s (i.e. perform worse) for speech-modulated noise maskers (see Spectral Invariance: Sensitivity to Bandpass Filtering) than the models that were trained on unfiltered data.

**Figure 8.**
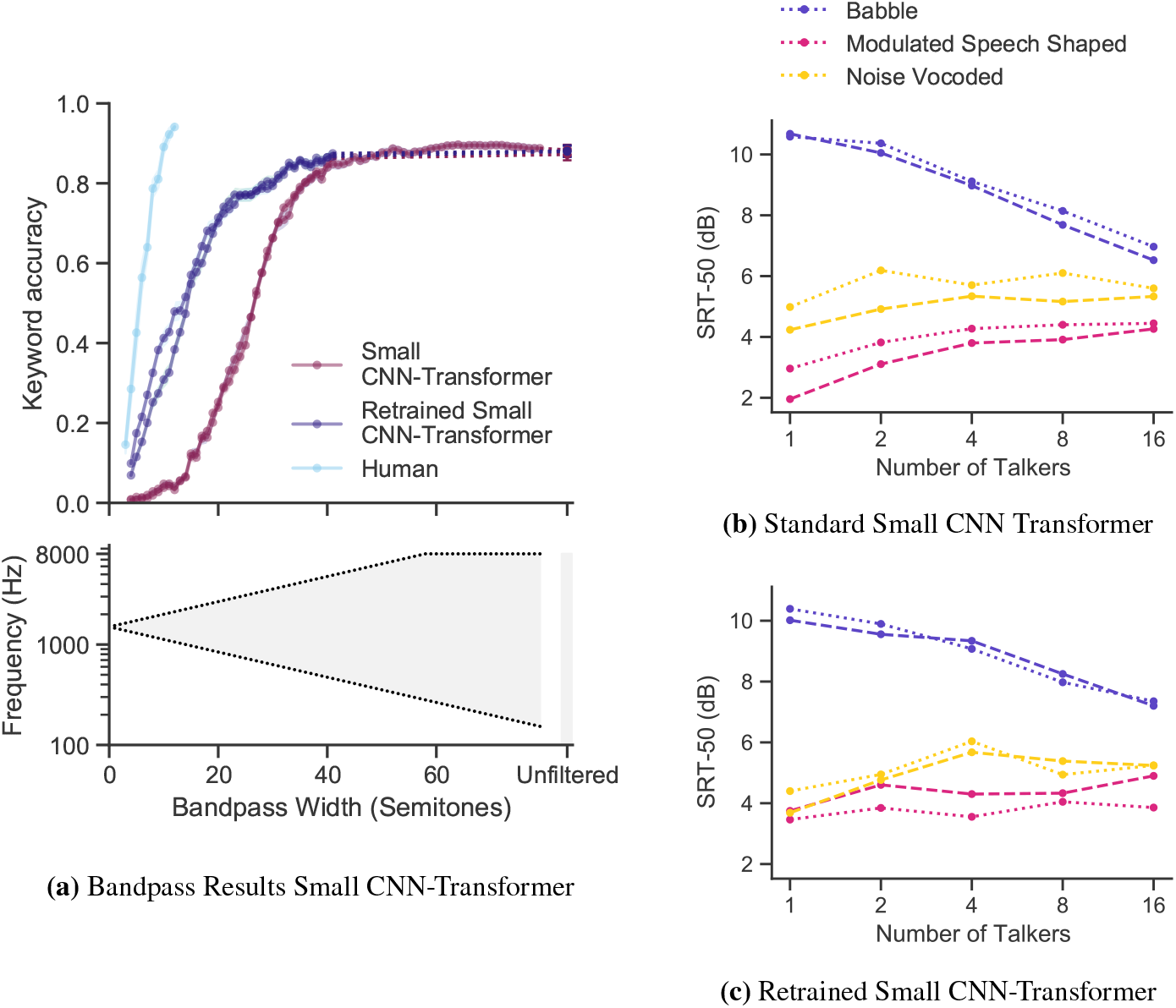
Performance of four smaller CNN-Transformer models, two of which were finetuned on bandpass filtered data, tested on the bandpass filter and masker periodicity experiment. (**a**) In the bandpass filter experiment, speech is bandpass filtered using a filter centered at 1500 Hz with varying widths (in semitones). Here we plot the keyword accuracy as a function of the bandpass width (in semitones) at which the target speech was filtered. Same data and procedure as for Figure 1a, human data from Warren et al. (2000). (**b-c**) Speech reception thresholds (SRTs) at 50% keyword accuracy for speech in three types of noise (babble, noise vocoded babble and speech-shaped noise modulated with babble envelope) with an increasing number of speakers. **b** displays the SRTs for the two small CNN-Transformers trained on unfiltered data whereas **c** shows the SRTs measured using two small CNN-Transformers that were trained on a mixture of normal and bandpass filtered data. SRT-50s were measured in the same way as for Figure 5.

When we are merely interested in creating an ASR system that performs well in a specific target task, increased performance in a single dimension is likely to be sufficient. A model can also be trained on data that was distorted in multiple dimensions to further increase robustness in a multi-task setting. However, our experiment suggests that such training may have unintended side effects. In particular, training on specific distortions may affect performance in currently unknown but nevertheless important dimensions. When the goal is to use ASR systems as a proxy for human listeners, these problems may be evaded by only training the model in realistic natural settings, such as a background of competing talkers. In the same way that human hearing is robust against certain distortions without much training, robustness against unnatural distortions may emerge in a candidate ASR system as a result of the way it has learned to account for natural noise. A test battery such as the one presented in this paper can be a good starting point for finding an ASR system that displays such consistent humanlike behaviour.

## 3 Discussion

We compared the performance of automatic speech recognition (ASR) systems and human listeners on a range of psychometric experiments. In general, ASR systems unsurprisingly performed worse than humans, particularly in the presence of noise. Where appropriate, we corrected for this by allowing a higher signal to noise ratio when testing the ASR systems. Even with this correction, none of the ASR systems performed overall similarly to humans in terms of which features and cues they relied on, and which distortions they were robust to. We conclude that they are not yet ready to be used as general purpose proxies for modelling human speech recognition, although they may be useful to ask more narrowly defined questions (which we discuss below).

The Wav2Vec 2.0 CNN-Transformer model, which is both the most recently introduced and model with most parameters of those being compared, was shown to be most robust against most distortions despite having been trained on less data. This model was most robust against bandpass filtering, spectral and temporal modulation filtering and various types of additive noise. The Kaldi DNN-HMM model, which had only slightly lower accuracy levels than the CNN-Transformer in quiet and undistorted listening conditions, tended to perform similar or worse in most dimensions. Mozilla’s DeepSpeech LSTM, which is the smallest model of those considered here, was much less robust than the two other systems.

The first three tests in our battery assess robustness to some common distortions found in communication systems: bandpass filtering (commonly applied in phone communication), peak clipping (a common distortion when a recording device is saturated) and centre clipping (as found in simple noise-suppression systems). We find that both the CNN-Transformer and the DNN-HMM closely follow human performance with regards to centre clipping, but none of the ASR systems come close to HSR performance with regards to bandpass filtering and peak clipping. Relative to the other ASR systems, the CNN-Transformer model displays the highest spectral invariance, whereas the LSTM model performs poorly in all three tasks.

The next set of tests investigated auditory features that have been commonly proposed to be important for speech processing in humans, namely spectral and temporal modulations, temporal fine structure (TFS) and periodicity. The tests showed that the relative sensitivity to spectral and temporal modulations were comparable between humans and ASR systems, but the use of TFS and periodicity differed considerably. The sensitivity to spectral and temporal modulations was consistent across ASR systems, with only a slightly higher importance of slow spectral and temporal modulations compared to humans. Disrupting TFS only affects intelligibility for normal-hearing humans in noisy conditions. In contrast, the ASR systems had only a tiny sensitivity to TFS, an effect that completely disappeared when notches of TFS were disrupted that did not span the whole frequency range. This experiment also further confirmed differences in spectral sensitivity between humans and ASR systems, as the removal of high frequency regions (above 2 kHz) hardly affects intelligibility for normal-hearing humans, but reduced speech in noise perception for all ASR systems. Lastly, the ASR systems appear to process periodicity information very differently from humans. Whereas humans perform poorly when vocoded speech is fully periodic (rather than mixed periodic as with natural speech), this effect was only observed for the CNN-Transformer. However, the CNN-Transformer performed better with noise-vocoded speech than with mixed periodic speech. Both the LSTM and DNN-HMM rely on the presence of periodicity and perform poorly in noise-vocoded speech. In stark contrast with humans, fully periodic vocoded speech does not negatively impact their performance compared to mixed-periodic speech.

The final set of tests tease out the factors involved in understanding speech in masking noise, including babble, modulated noise and periodic maskers. The CNN-Transformer model is most similar to humans and can make use of both glimpsing opportunities and masker periodicity to the same extent as humans, although it does suffer more from informational masking from noise-vocoded speech. The DNN-HMM model is able to make use of glimpsing opportunities in speech-shaped modulated noise, but this ability does not extend to 10 Hz noise modulations. The LSTM model shows no benefit for masker periodicity or modulations. No benefit of having only a single competing talker was observed for any of the ASR systems, although this is likely the result of the fact that the ASR systems could not be instructed to attend to a particular speaker, whereas the human listeners were. It would be interesting to see how the performance in a competing talker experiment changes for models that are able to focus on a specific speaker.

Altogether, these results highlight the differences between ASR systems and humans. Even when they achieve similar performance in quiet listening conditions, not only are the ASR systems much less robust than humans to common distortions such as bandpass filtering and peak clipping, but all models perform consistently worse than humans under the same SNR conditions. As none of these models were trained in noisy conditions, these findings are not surprising. However, we also showed that fine tuning models with certain distortions may negatively affect performance in other dimensions. Considering that the CNN-Transformer model is the most robust against distortions despite being trained on the least varied data, we conclude that it is not sufficient to simply increase the types of data it is trained on, but that further developments of the model and training procedure are required. Our findings are consistent with other work comparing human and automatic speech recognition (Adolfi et al., 2022), and work that points to the robustness to mismatched train and test domains for pre-trained acoustic representations such as those used in the Wav2Vec 2.0 CNN-Transformer (Ma et al., 2020).

Looking to the future, designing ASR systems that rely on similar features of speech as humans may allow them to perform better in noisy conditions they have never been exposed to. Our results point to some features that may be important. For example, none of the ASR systems presented here are able to exploit TFS information. In the case of the LSTM and DNN-HMM, this is the result of the use of MFCC features as input. The end-to-end ASR systems, however, should in principle be able to utilise TFS information. Other dimensions in which the ASR systems differed significantly from humans are resistance to peak clipping distortions and the use of periodicity in the target speech.

Despite these differences, the CNN-Transformer model tested here may already be similar enough to humans to be used to generate some narrow hypotheses about human hearing. Particularly, the similarities in sensitivity to glimpsing and masker periodicity may be used to identify previously unknown auditory features or mechanisms. For example, Steinmetzger and Rosen (2015) suggested a range of mechanisms and features such as harmonic cancellation, modulations and spectral glimpsing may underlie the periodicity benefit observed for humans. By applying explainable machine learning techniques to understand the mechanisms that underlie the masker periodicity benefit in the CNN-Transformer model, the relative contribution of these different mechanisms may be better understood, or other mechanisms could be identified.

It may also be fruitful to investigate why the CNN-Transformer model performs more similarly to humans than the other models. An immediate possibility is the unsupervised pretraining of the CNN-Transformer. In contrast to the other two models, the CNN-Transformer model does not use MFCC features but instead employs unsupervised pretraining to automatically extract acoustic features from speech data. This results in a rich speech representation that captures a wide range of phonetic information (Ma et al., 2020) as well as a surprising amount of higher level linguistic structure (Shah et al., 2021). Other pretrained acoustic representations have shown similar robustness to mismatched train and test data (Ma et al., 2020), which suggests that unsupervised, task-independent pretraining is an important step towards creating more humanlike ASR systems.

Finally, we have released all our code as an easily extensible open source toolbox, HumanlikeHearing. The toolbox can be used to find new directions for ASR research by exposing differences in the use of, for example, TFS and target periodicity information. It can also be used to find similarities between ASR systems and humans in order to identify the best ASR systems that could be used as a proxy for human listeners for hypothesis generation.

## 4 Methods and Materials

### 4.1 Speech Dataset

The main dataset used in our experiment is the ARU speech corpus (Hopkins et al., 2019), a freely available dataset which consists of recordings of the IEEE (Harvard) sentences (IEEE, 1969) spoken by twelve adult native British English speakers in anechoic conditions. The IEEE sentences are commonly used in auditory tasks because they are designed to be phonetically balanced and use specific phonemes at the same frequency as they appear in English. However, they do not contain a strong semantic signal: the sentences are gramatically correct but make little to no sense. All sentences are of approximately the same length and are short so that they can easily be repeated by human participants. All of the experiments measure performance as percentage of keywords correct. To automatically extract keywords from any data set, part-of-speech (POS) tags were determined using the Natural Language Toolkit (NLTK, Loper and Bird (2002)). Then, a heuristic was applied in which groups such as nouns and verbs were included as keywords, whilst ignoring auxiliary verbs and forms of *to be*. For the IEEE dataset, this approach led to the selection of approximately five keywords per sentence. By selecting kewords automatically, the Humanlikehearing toolbox can easily be extended to other data sets.

The three ASR models were trained on publicly available speech data. The CNN-Transformer was pre-trained on the LibriSpeech dataset, a corpus of approximately 1000 h of read English speech derived from audiobooks of the LibriVox project (Panayotov et al., 2015) in 16 kHz. It was fine-tuned on a subset of 100 h of labeled clean speech. The DNN-HMM and LSTM models use the full LibriSpeech dataset for training together with other publicly available English training data, leading to a total of 3000 h and 3800 h for DNN-HMM and LSTM, respectively. To the best of our knowledge, none of these publicly available datasets incorporate IEEE sentences.

### 4.2 Sound Level Normalisation

Before applying any of the tests in the test battery, it is important to ensure all sounds are normalised in the way the ASR system expects. In the toolbox, sound files in a range of formats (such as WAV or FLAC files) are loaded as floating point values ranging mostly between -1 and 1. Under the assumption that these values represent the sound pressure in Pa, the root mean squared (RMS) and the sound level (SPL) in dB is computed as follows:

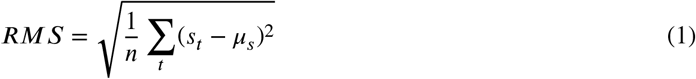

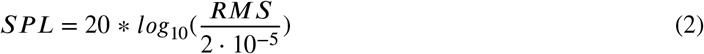

Here *s*_*t*_ represents the sample of a signal *s* at time *t, n* reflects the total number of samples in s and *µ*_*s*_ is the average value of all samples in *s*. The factor 2 · 10^−5^ reflects the reference pressure of the smallest sound humans can hear in Pa. In addition to this basic RMS-based sound level, the toolbox offers more advanced ways of estimating sound levels. Firstly, A-weighting can be applied to allow for the fact that the perceived loudness of a sound given a certain acoustic pressure varies depending on the frequency of the sound. Furthermore, a sound level computation for speech signals has been implemented that adjusts the speech level based on a speech activity detector. This implementation, which follows the IEEE P56 recommendation, mitigates the fact that in the normal RMS computation moments of silence in a speech segment skew the SPL to be lower than it actually is.

Based on the computed sound levels, a preferred signal to noise ratio (SNR) can be achieved by either varying the noise level or the speech level to achieve the required SNR. The level is adjusted by multiplying each sample in the sound with a gain that is computed as follows:

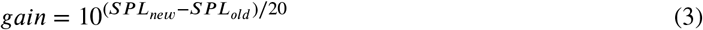

Here, *SP L*_*oid*_ is the current sound level and *SP L*_*new*_ is the required sound level. In most of our experiments, the speech levels are held constant at 65 dB SPL, and the noise levels are varied accordingly. In humans, setting the level to around 65 dB SPL will result in a comfortable loudness. To determine the preferred loudness of the ASR system that is being tested, the toolbox includes a ‘normalisation test’ that measures ASR performance for clean speech as a function of sound level in dB SPL. Applying this test shows that the CNN-Transformer (which normalises input sounds) performs well regardless of sound level (Figure 9), whereas the LSTM and DNN-HMM only perform well within the ranges of 50 dB to 80 dB SPL and 30 dB to 90 dB SPL, respectively.

**Figure 9.**
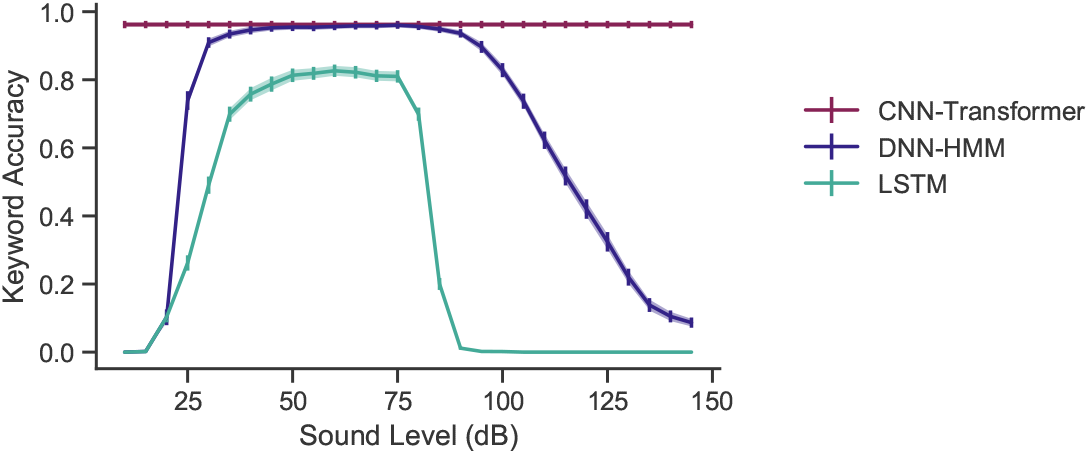
Sound level sensitivity of the three different ASR systems tested on the first 10 lists (100 sentences total) of the IEEE dataset for the three different ASR systems. Performance is measured as percentage of keywords correct with approximately five keywords per sentence. Errorbars denote the standard error of the mean.

### 4.3 Speech Reception Threshold Determination

The performance of listeners in a speech in noise experiment is often summarised using the Speech Reception Threshold (SRT), which refers to the SNR at which a certain level of accuracy (usually 50% or 70.7%) is achieved. The SRT is determined by fitting a psychometric function (usually of sigmoidal shape) to a set of SNR values and accuracy pairs. To fit the psychometric function to a given set of samples, we used psignifit 4, a software package that implements the maximum likelihood procedure as described in Wichmann and Hill (2001). When estimating SRTs, we leave the lapse rate (which sets an upper limit to performance) as a free parameter.

To reduce the number of trials per condition for humans, data points are usually collected through an adaptive procedure. For ASR systems, which unlike humans do not display attention fatigue or sentence memorisation, it is possible to measure the psychometric curves more precisely through a grid search that tests every sentence on a range of SNRs. We refer to the latter as the static SRT procedure. Both the adaptive and static procedures are implemented in the toolbox.

In adaptive procedures, the SNR is first set to a value at which performance is expected to be good. This point can be estimated adaptively using the first sentence only by starting at a very low SNR, after which the SNR is increased until the sentence is exactly identified. However, as it is not guaranteed that an ASR system can achieve 100% performance, the maximum accuracy is set to the accuracy in quiet in the toolbox. For subsequent sentences, the SNR is either increased (or decreased) depending on whether performance was worse than (or better than) the target threshold. The step size of the SNR changes is usually reduced after a given number of ‘reversals’. For example, Rosen et al. (2013) reduced the step size from 4 dB to 2 dB once the direction of change of the SNR had reversed twice. As a simpler alternative to Maximum Likelihood Estimation (MLE), ‘staircase’ sampling methods are sometimes used (Leek, 2001). Here, a response is assumed to be either correct or incorrect (positive or negative). To target SRT-50, a one-up one-down procedure is used, where a positive trial results in a decrease of SNR and a negative trial results in an increase. To target SRT-70.7, a two-down, one-up procedure is used, where two positive responses are required to result in a lowering of the SNR. This two-down, one-up procedure was used in the temporal fine structure experiment described in Section Temporal Fine Structure (Hopkins and Moore, 2010). From this data, the SRT is sometimes estimated by taking the mean of all the SNR values at the last few reversals. A disadvantage of such staircase methods is that the response needs to be reduced to a correct/incorrect response. This is a reasonable assumption in closed-set speech recognition tasks, such as is the case in the coordinate response measure (CRM) dataset used by Hopkins and Moore (2010), but less applicable to the more complex IEEE sentences used in this work.

In the static SRT procedure, a reasonable range of SNR values can be determined through a pilot study using a small set of sentences. In contrast to the adaptive procedure, which targets SNR values around the SRT, the static procedure obtains data points over the whole SNR range and can thus be used to determine the SRT at any performance level. However, as it collects more data points it is much slower than the adaptive procedure. All plots in the paper are measured using the static approach. However, provided performance in quiet was higher than the SRT accuracy level, the adaptive sampling method gave very similar results.

### 4.4 Signal Processing and Procedures

#### Spectral Invariance: Sensitivity to Bandpass Filtering

To simulate the experiment presented in Warren et al. (2000), each sentence was filtered using a bandpass filter centered at 1500 Hz. The 2000-order finite impulse response (FIR) filter used has a steep slope of 1000 dB/octave. This steep slope removes the presence of transition bands, which may otherwise leak information from other frequency regions. In Warren et al. (2000), the bandwidth varies in 1-semitone steps with values ranging from 3 semitone up to 12 semitones, but this range was extended to 80 semitones to ensure all ASR systems reached ceiling-level performance.

Instead of the CID Everyday Sentences corpus (100 sentences, male speaker) used in Warren et al. (2000), which to the best of our knowledge is not freely available, we used the IEEE sentence lists in our experiment. The IEEE sentences are of similar length to the CID sentences, and in both corpora each list of 10 sentences contains 50 keywords. However, CID sentences are more semantically meaningful than IEEE sentences, which makes it easier for human listeners to guess unintelligble words. A lack of semantic context in bandpass filtered speech has been reported to lead to an accuracy reduction of around 20% (Stickney and Assmann, 2001). When considering words in isolation, i.e. without grammatical context, performance drops a further 23%. The intelligibility of bandpass filtered IEEE sentences in humans is thus expected to be lower than that of the CID sentences. However, since it is unlikely that ASR systems can utilise semantic context to the same extent as humans can, we expect that ASR performance for IEEE and CID sentences would be relatively similar.

#### Peak and Centre Clipping

To study the robustness of the ASR systems to both peak and centre clipping, the experimental setup presented in Kates and Arehart (2005) was implemented. In this study, clipping distortions are parameterized by a threshold *t* which ranges between 0 and 1. To determine the clipping value *c* associated with this threshold, silence intervals at the start and end of the sentences are removed and a histogram of the magnitudes of the signal samples is computed. The clipping value *c* is set to the percentage *t* of the cumulative magnitude histogram for the sentence.

Given the clipping value *c*, the peak clipping operation is given by:

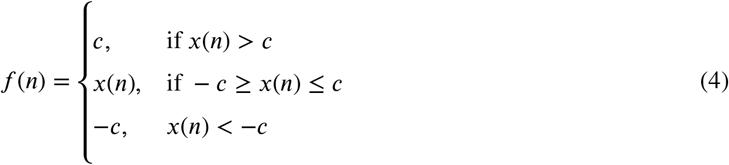

Here, *x*(*n*) is the *n*th sample of the speech signal, and *f* (*n*) the distorted output sample. The centre clipping operation is defined as follows:

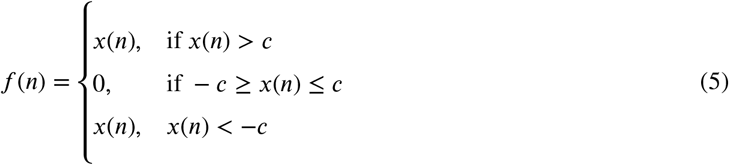

The distorted sentences are then readjusted to a sound pressure level of 65 dB. As in Kates and Arehart (2005), the peak clipping thresholds were set to 0%, 50%, 75%, 90%, 95%, 98%, 99% and 100% of the cumulative magnitude histogram of each sentence. The centre clipping thresholds were set to 0%, 50%, 70%, 80%, 90%, 95% and 98%. In Kates and Arehart (2005), each of the thirteen listeners was presented with approximately 10 sentences for each threshold condition, giving a total of 130 HINT sentences per threshold condition. In the present study, 100 IEEE sentences were used per condition. The IEEE and HINT sentences are of similar length, but the IEEE sentences provide less semantic context. This difference should not affect the results too much, as intelligility rates measured for clipped isolated words without grammatical or semantic context lead to similar intelligibility rates to those reported in Kates and Arehart (2005), as shown in (Licklider and Pollack, 1948).

#### Spectral and Temporal Modulations

The temporal or spectral modulations of a signal can be extracted by computing the Modulation Power Spectrum (MPS), a 2D representation of a signal in which each axis represents either spectral or temporal modulations. To remove specific modulations from a target speech signal, the signal is transformed into the MPS, the desired modulations are set to zero, and the MPS is inverted back into speech. Following the procedure described in Elliott and Theunissen (2009), the MPS is derived by first computing a spectrogram using Gaussian windows on a linear frequency scale (the choice of a linear rather than log-spaced spectogram is why results are reported in cycles/kHz rather than octaves/kHz). The modulation power and phase spectrum are obtained from the log of the spectrogram using the 2D Fast Fourier Transform (FFT). The required modulations are set to zero in the power spectrum and randomised in the phase spectrum. To invert the filtered modulation spectrum back into a sound, the inverse 2D FFT is applied to obtain the filtered log spectrogram. This spectrogram is exponantiated and inverted using the Griffin-Lim iterative spectrogram inversion algorithm (Griffin and Lim, 1984) to obtain the filtered speech.

To assess the importance of specific spectral and temporal regions, we reproduced the notch experiment presented in Elliott and Theunissen (2009). In this experiment, speech filtered with a total of ten temporal and nine spectral modulations was embededded in Gaussian white noise with 2 dB SNR. In about half of the notches, the ‘core’ region (set to spectral modulations up to 3.75 cycles/kHz and temporal modulations up to 7.75 Hz) were removed before removing the notch. Performance was also measured against a control condition (spectrogram inversion without modulation filtering) and the core region (without modulation filtering). Each of the 17 listeners was presented with 100 sentences that were randomly assigned to a condition. Given 21 conditions (nine spectral, ten temporal notches, control and core), an average of 81 sentences total (five per listener) were presented per condition. Our results are based on 100 sentences per condition. Stimuli in Elliott and Theunissen (2009) are taken from the soundtrack of the Iowa Audiovisual Speech Perception videotape, which are of similar length but provide less semantic context than the IEEE sentences used here. Furthermore, to control for the poor robustness against white noise of the ASR systems, an additional noise condition was tested in which the SNR was set to the lowest SNR at which the ASR system achieved roughly ceiling performance (10 dB, 15 dB and 25 dB for the CNN-Transformer, DNN-HMM and LSTM, respectively). In Elliott and Theunissen (2009), SNRs were set relative to a constant noise level of 65 dB. To prevent the overall sound level from exceeding the preferred ranges of the ASR systems when testing at high SNR levels (see Sound Level Normalisation for details), we chose to set SNRs relative to a constant speech level of 65 dB.

#### Temporal Fine Structure

To investigate the usage of temporal fine structure in the ASR systems, we implemented two experiments described in Hopkins and Moore (2010) in which the use of TFS information is investigated in a cumulative as well as region-specific manner. Both experiments use a vocoding paradigm in which the speech is filtered into a range of frequency bands. Specifically, sounds are processed using a tone vocoder with 30 channels on an Equivalent Rectangular Bandwidth (ERB) scale (Glasberg and Moore, 1990) from 100 Hz to 8 kHz. The linear-phase FIR filters had a variable order, chosen such that the transition bands of each filter had similar slopes on a logarithmic frequency scale. Each filter was designed to have a response of −6 dB relative to the peak response at the frequencies at which its response intersected with the responses of two adjacent filters. The channels were divided into five regions, each spanning 6 ERB_*N*_. To retain both TFS and envelope information, the channel was left unchanged. To distort only TFS information but retain the channel envelope, a sine wave of the channel centre frequency was modulated with the channel envelope, which was extracted using a Hilbert transform. The modulated tone was then filtered again with the channel bandpass filter to ensure side bands are removed.

In the first experiment, TFS information was added successively either starting from the low frequency region (*TFS-low*) or high frequency region (*TFS-high*). In the second experiment, all channels are tone-vocoded except for one region. The channels in this region are processed in one of two ways. In the *+* condition, the channels remain unprocessed, thus retaining intact TFS. In the *-* condition, the channels are set to zero, thus removing both envelope and TFS information. Performance is also measured in the *allvoc* condition, in which all channels are tone-vocoded. The benefit of the envelope in a specific region is computed as the difference between the SRT found in the *allvoc* and *-* condition, whereas the combined benefit of the envelope and TFS is the difference between the the *-* and *+* condition.

In our experiments, the SRT-70s of ASR systems were measured five times using 20 IEEE sentences (100 different sentences total) in six-speaker babble noise. We used a static SRT determination technique with SRTs ranging from −4 dB to 24 dB in 2 dB steps. As in Hopkins and Moore (2010), the target speech level was fixed at 65 dB.

To ensure consistency throughout this work, performance in our experiments is measured agaist the IEEE dataset rather than the Coordinate Response Measure (CRM) set used in Hopkins and Moore (2009). In the CRM corpus, sentences consist of only three words that follow a common structure (*“Ready [call sign] go to [colour] [number] now*.*”*) and are selected from eight call signs, four colours and eight numbers. In such a closed-set task, it is much easier to achieve higher SRTs. This is particularly the case for humans, as they can be instructed to only pick words from the available set. For completeness, we include results on the CRM corpus in the appendix (see Temporal Fine Structure on CRM corpus).

Another way in which our version of the experiment differs is the choice of masker. We used a babble noise of six talkers rather than a single talker. This was done to account for the fact that the ASR systems used here cannot attend to a specific speaker and, as shown in Competing Talker Backgrounds, are much worse at recognising speech in a single competing talker than humans are. The use of different target and masker datasets is unlikely to lead to a qualitative difference in results. In a comparable experiment, SRT-50 curves for the *TFS-low* condition were qualitatively similar despite being measured using IEEE sentences in speech-shaped noise (Hopkins and Moore, 2009).

#### Target Periodicity

In the target periodicity experiment, the intelligibility of three types of speech was investigated: aperiodic (noise-vocoded) speech, speech with a natural amount of source periodicity and fully periodic speech following the procedure described in Steinmetzger and Rosen (2015). The *F*_0_ contours of the speech signal were extracted using PRAAT with a sampling rate of 100 Hz (Boersma, 2021). In our toolbox, the Python module Parselmouth is used to interface with PRAAT software (Jadoul et al., 2018). To obtain the vocoded speech, the original recordings were first bandpass filtered into 6, 7, 8, 10, 12, 16 or 24 bands using a sixth order Butterworth IIR filter. Filters were spaced in a a range from 100 Hz to 11 kHz following equal basilar membrane distance (Greenwood, 1990). To extract the envelope, the output of each band was full-wave rectified and lowpass filtered at 30 Hz using a fourth-order Butterworth filter. For the noise-vocoded speech, the envelope was multiplied by a wide-band noise carrier. For Dudley vocoded speech, which has a natural amount of source periodicity, and fully periodic speech, the envelope of the voiced segments of speech was multiplied with a pulse train following the natural *F*_0_ contour, and the segments of unvoiced speech were multiplied with a wide band noise carrier. The periodic speech was generated using pulse trains only. Here, additional *F*_0_ contours were created by interpolation through unvoiced sections and periods of silence using piecewise cubic Hermite interpolation of a logarithmic frequency scale. The start and end points of each contour were anchored to the median frequency of the sentence. For all three vocoding types, the side bands were removed by filtering the modulated noise and/or pulse train using the channel bandpass filter. The RMS level of the channel outputs was then adjusted to match the original level in that band, after which all band filters were summed. The final waveforms were then low-pass filtered at 10 kHz.

In Steinmetzger and Rosen (2015) a second set of non-periodic, mixed periodic and fully periodic speech was generated using TANDEM-STRAIGHT (Kawahara et al., 2008). This method does not rely on bandpass filtering but produces very natural-sounding speech with a mixed source excitation that can be adapted to produce fully aperiodic or fully periodic speech. Using TANDEM-STRAIGHT, non-periodic speech was generated by keeping the default settings but fixing the *F*_0_ to 0 Hz throughout. For Dudley vocoded speech, to minimise the level of the aperiodic component the values of the sigmoid parameter in the source estimation routine were fixed to 1 and −40. For *F*_0_-vocoded speech the interpolated *F*_0_ contours were used as input for the source extraction routine.

Since Steinmetzger and Rosen (2015) made their material publicly available, the results reported in this section are based on their exact stimuli. The toolbox contains an implementation that leads to very similar results for the vocoded speech. However, since TANDEM-STRAIGHT is patent-protected software it is not included in the toolbox. The stimuli are recordings of ten lists of IEEE sentences (100 sentences total) spoken by a British male. A total of 220 sentences (20 sentences per 11 participants) were presented for each condition (a particular type of vocoding with a given number of channels). Our results are based on 100 sentences per condition (i.e. all 10 IEEE lists). In Steinmetzger and Rosen (2015), the level of the target was fixed at about 80 dB SPL, whereas here it is fixed at 65 dB SPL based on the normalisation test results of the ASR systems.

#### Competing Talker Backgrounds

In the competing talker background experiments, the three ASR systems were tested using the three masker conditions from Rosen et al. (2013). These conditions include speech babble, noise-vocoded babble and speech-shaped noise that was modulated using the envelope of the babble noise. The basis for their noise maskers were male talkers from the EUROM database of English speech, a dataset in which each speaker reads five or six passage sentences. Of these, pauses of more than 100 ms were deleted, resulting in sound files of approximately 21 s per speaker without significant pauses. In the HumanlikeHearing toolbox, segments of similar length are created by combining sentences spoken by male speakers from the IEEE ARU database. The sound files were normalised to a common RMS values. Speech babble was created by randomly selecting one of the sixteen background talkers. As the ARU dataset only contains 6 male talkers, some talkers are sampled multiple times in the toolbox. The other talker conditions were creating by randomly selecting a talker and adding it to the talker(s) already present in the previously constructed condition. The spectrum of each babble was equalized to the long-term average spectrum of the 16-talker speech babble. To create noise-vocoded babble, each of the five babbles was filtered into 12 bands using the method described in Target Periodicity. To create speech-shaped modulated noise, the envelope of each babble was extracted by full-wave rectification and low-pass filtering at 30 Hz. The envelope was multiplied by a broad-band noise that was equalised to the long-term average spectrum of the 16-talker babble.

The results presented here use the same material as in the original experiment of Rosen et al. (2013), but the toolbox also incorporates an implementation that leads to similar results. The target speech consists of the first 10 lists of the IEEE dataset. In Rosen et al. (2013), SRT-50s were obtained using an adaptive approach. The results reported here were measured using the static SRT method instead, where data obtained over a range of SNRs were fitted with a logistic function using a Bayesian optimisation paradigm. Specifically, SRT-50s were estimated from scores at SNRs measured ranging from −6 dB to 33 dB in 3 dB steps. Rosen et al. (2013) presented 320 sentences per condition (20 sentences for each of the 16 listeners). Our experiments are based on the results of all 100 sentences. The noise masker was randomly selected from the 21 s masker. In Rosen et al. (2013), the noise level was fixed at 70 dB SPL over a frequency range of 100 Hz to 5 kHz. To obtain a given SNR, the target speech level was adjusted. In our experiments, the target speech remained fixed at 65 dB SPL throughout.

#### Masker Modulations and Periodicity

To investigate the effects of masker modulations and periodicity, we followed the second and third experiments described in Steinmetzger and Rosen (2015). The target speech was the same as described in Target Periodicity, i.e. non-periodic (noise-vocoded), mixed-periodic (Dudley vocoded), fully periodic as well as unfiltered speech. Two types of maskers, a speech-shaped noise masker and a harmonic complex, were used. The noise masker was a 24 s passage of white noise that was filtered to follow the long term average speech spectrum (LTAS) of the unprocessed target speech. The LTAS was computed by taking the power spectral density of the concatenated waveforms using Welch’s method (with a window size of 512 samples, 50% overlap and FFT length 512 samples), followed by 1-octave smoothing. The harmonic complex maskers were based on *F*_0_ contours extracted from recordings in the EUROM database. In the toolbox, they are extracted from concatenated IEEE sentences. The contours were extracted using PRAAT and interpolated through unvoiced and silent periods using a piecewise cubic Hermite interpolation on a logarithmic frequency scale. The waveforms were synthesized on a period-by-period basis using the Liljencrants-Fant model (Fant et al., 1985), which closely approximates a typical adult male glottal pulse. In the toolbox, the glottal pulse model incorporated in PRAAT is used instead. Using the same filtering procedure as for the noise maskers, the harmonic complex maskers were matched in spectrum to the LTAS. Both the noise and harmonic complex maskers were presented either as is or were sinusoidally amplitude modulated at a rate of 10 Hz with a modulation depth of 100%. During the experiment, the maskers were randomly selected from the 21 s segment.

As in Target Periodicity, the results reported here are based on the material made publicly available by Steinmetzger and Rosen (2015). In their study, a total of 240 sentences (20 sentences per 11 participants) were presented for each condition (a combination of one of the three vocoded targets with a certain spectral resolution or unfiltered speech together with one of the four maskers). Here, each condition is tested using 100 sentences (all available IEEE sentences). In Steinmetzger and Rosen (2015), each vocoded target was tested under three (for noise vocoded and Dudley vocoded) or four degrees (fully periodic vocoded) of spectral resolution. The spectral resolution for each vocoder type was chosen to match performance in quiet to around 70%, 90% or ceiling-level. In our experiments, the SRT-50 was measured for all levels of spectral resolution in which performance in quiet was above 50%. In Steinmetzger and Rosen (2015), the level of the target and masker together was fixed at about 80 dB SPL, whereas in the results presented here the target speech was fixed at 65 dB. Steinmetzger and Rosen (2015) determined SRT-50 using an adaptive procedure, whereas the results reported here are based on the static SRT method (see Speech Reception Threshold Determination).

## 5 Acknowledgments

This work was partly supported by a Titan Xp donated by the NVIDIA Corporation, Royal Society grant RG170298 and Engineering and Physical Sciences Research Council grant number EP/L016737/1.

## A Details of ASR Systems

Three freely available ASR models are analysed in this work: the Kaldi nnet3 chain model (trained as part of the Kaldi Active Grammar project), DeepSpeech (trained as part of Mozilla’s DeepSpeech project) and Wav2Vec 2.0 (trained as part of Facebook’s fairseq toolbox). We used out-of-the-box models that have been made freely available by their developers using a default setup. Specifically, the models used were as follows:

1. Kaldi nnet3 chain model trained by David Zurow, named *vosk-model-en-us-daanzu-20200905-lgraph*. Available for download at https://alphacephei.com/vosk/models
2. Mozilla’s DeepSpeech 0.6.1 release, available for download at https://deepspeech.readthedocs.io/en/v0.6.1
3. Fairseq’s Wav2Vec 2.0 Large 960 model, available for download at https://github.com/pytorch/fairseq

Each of these models relies on a different class of commonly used ASR architectures, which are described in more detail below.

Vosk’s Kaldi nnet3 chain model is a **DNN-HMM** hybrid model, a type of model that combines deep neural networks (DNN) with Hidden Markov Models (HMM) (Povey et al., 2018). The model takes as input Mel-Frequency Cepstrum Coefficients (MFCC) features and i-vectors. MFCCs are commonly used ASR input features that concisely describe the overall shape of a spectral envelope of a sound wave. The DNN-HMM model extracts 40 MFCCs from a spectrogram ranging from 20 Hz to 7600 Hz, with windows of 25 ms shifted by 10 ms each time. I-vectors are latent variables that model speaker-specific characteristics of speech and can be obtained through factor analysis (Dehak et al., 2010). The MFCC features and i-vectors are fed into a 15-layer factorised Time Delay Neural Network (TDNN-F) (Povey et al., 2018). The layers of a TDNN are designed to classify patterns with shift-invariance, which means the classifier does not require explicit segmentation. Shift-invariance is achieved by averaging the backpropagation update over time-shifted copies of the network across the temporal dimension. Another feature of TDNNs is that each layer receives not only the output of the previous layer at the current time step, but receives a contextual window of outputs from the layer below. This allows the TDNN to model the temporal context. To efficiently model long temporal contexts, the Kaldi nnet3 chain model uses a subsampling technique in which hidden activations at only a few time steps are computed at each level, while ensuring that information from all time steps in the input context is processed by the network (Peddinti et al., 2015). The TDNN-F applies SVD-like model reduction at every layer, which reduces model size by effectively applying a bottleneck to each layer. In the DNN-HMM model, layers of 1536 hidden neurons are reduced to a bottleneck layer of 160 nodes. The model is trained using SpecAugment (Park et al., 2019), a data augmentation technique, and an end-to-end variant of the lattice-free maximum mutual information (LF-MMI) criterion (Povey et al., 2016).

Mozilla’s DeepSpeech model is a Long-Short-Term-Memory (**LSTM**) neural network, which is a specific type of neural network that can model long temporal relationships in the input data. Mozilla’s DeepSpeech is a variant of the original DeepSpeech model (Hannun et al., 2014), which used simpler recurrent neural network (RNN) units. In contrast to the original DeepSpeech model, which takes the raw waveform as input, the input to Mozilla’s DeepSpeech model consists of 26 MFCCs that have been extracted from a spectrogram in a range of 20 Hz to 4 kHz with a window length of 32 ms and a stride of 20 ms. One set of MFCCs is referred to as a frame and reflects the spectral content over the 20 ms window. The model architecture consists of six layers, all of which are standard feed-forward layers except for the fourth layer, which consists of recurrent LSTM units. The last layer is a softmax layer. The outputs of the model are letters (non-capitalised) and the model is trained using the Connectionist Temporal Classification (CTC) criterion (Graves et al., 2006).

The last model considered is Facebook fairseq’s Wav2Vec 2.0, which consists of a convolutional neural network (CNN) and a Transformer model (**CNN-Transformer**, Baevski et al. 2020). In contrast to the other two ASR systems, the CNN-Transformer takes the raw audio as input. To extract relevant acoustic features from the audio, the CNN-Transformer is initially ‘pre-trained’ in a self-supervised fashion. First, raw audio data is encoded into a latent speech representation by a seven-layer CNN that functions as a feature encoder. These representations are fed into a Transformer, which is used to build representations that capture information from the entire sequence. The model used here consists of 24 transformer blocks with model dimension 1024, inner dimension 4096 and 16 attention heads. To train the CNN-Transformer, the output of the feature encoder (i.e. the CNN component) is discretized. These discrete units are then used to represent targets in a self-supervised objective, which requires the model to identify the correct quantized latent audio representation in a set of distractors. After pre-training, the model is fine-tuned for speech recognition by adding a randomly initialised linear projection on top of the context network into classes that represent the vocabulary (e.g. letters) of the task. The whole model is then optimized for speech recognition by minimizing the CTC loss.

Some basic limits on the spectral and temporal components of speech that each of these models can utilise can be derived from their inputs and architectures. For example, as all models are trained on sounds sampled at 16 kHz, frequency information of at most 8 kHz is present in the input. The LSTM is limited to 4 kHz, as the MFCCs are derived from a spectrogram limited to that frequency. In the DNN-HMM and LSTM architecture, spectral information is initially captured by the MFCC representation, in which much of the temporal fine structure (TFS) is lost. In the CNN-Transformer model, spectral information is extracted by the CNN layers, which effectively function as Finite Impulse Response (FIR) filters. The latent speech representations generated by the encoder of the CNN-Transformer model have a window size and stride of approximately 25 ms and 20 ms, respectively, which is comparable to the window and stride of the MFCC features of the DNN-HMM (25 ms and 10 ms) and LSTM model (32 ms and 20 ms).

To model long temporal relationships between frames, each model uses a different strategy. In the TDNN-F architecture used by the DNN-HMM, each layer at time *t* receives the outputs of the previous layer at times between [*t* − *s, t* + *s*], where *s* denotes the ‘time stride’ of that layer. Given the time strides (1, 1, 1, 0, 3, 3, 3, 3, 3, 3, 3, 3, 3, 3) of the 14 TDNN-F layers in the DNN-HMM, the output layer has access to temporal information from 67 MFCC frames (685 ms). In the LSTM model, the input of the model at any given timestep consists not only of the MFCCs for that specific frame, but also those of the nine preceding and following frames. The model input thus covers a range of 19 MFCC frames (392 ms). Additionally, the LSTM layers can retain information over multiple timesteps, allowing the model to encode longer temporal relationships. In the CNN-Transformer model, temporal information between latent speech representations is handled by the Transformer architecture, which uses an attention mechanism to relate inputs from different timesteps. In theory, transformer models can span temporal relationships of infinite duration, but in practice this CNN-Transformer model was trained using sequences of at most 20 s.

## B Temporal Fine Structure on CRM corpus

As discussed in Temporal Fine Structure, the CRM corpus used in Hopkins and Moore (2010) is a closed-set corpus in which sentences have a common structure (*“Ready [call sign] go to [colour] [number] now*.*”*). The human participants in this experiment were instructed to pick one out of eight call signs, one out of four colours and one out of eight numbers. This makes the task much easier than recognising sentences from the IEEE dataset, which contains five keywords per sentence that are not selected from a predefined subset and are not strongly semantically related. To ensure that our results were not task-dependent, we repeated the experiment on the CRM corpus. To mimic the closed-set task in a model-agnostic manner, we replaced each predicted transcription with the most similar valid CMR sentence. This most similar sentence was determined by computing the character error rate (measured as the character-level Levenshtein distance) between the predicted transcriptions and all 256 (8 call signs, 4 colors, 8 numbers) possible sentences and selecting the sentence with the lowest error rate. As in Hopkins and Moore (2010), we then reduced the accuracy to a binary correct/incorrect that indicates whether or not the correct sentence was selected. We used the static SRT estimation procedure (see Speech Reception Threshold Determination). As shown in Appendix 2 Figure 1, the use of the simpler task results in higher SRTs, but leads to qualitatively similar performance. In particular, the cumulative TFS benefit is still not as high as it is in humans, and no significant TFS benefit is observed in the second notch experiment. However, we do note some small differences in the different spectral regions in the second experiment. In particular, the 4th spectral region appears less important for the LSTM than it was for IEEE data, although there is also a much higher standard error of the mean.

**Appendix 2 Figure 1.**
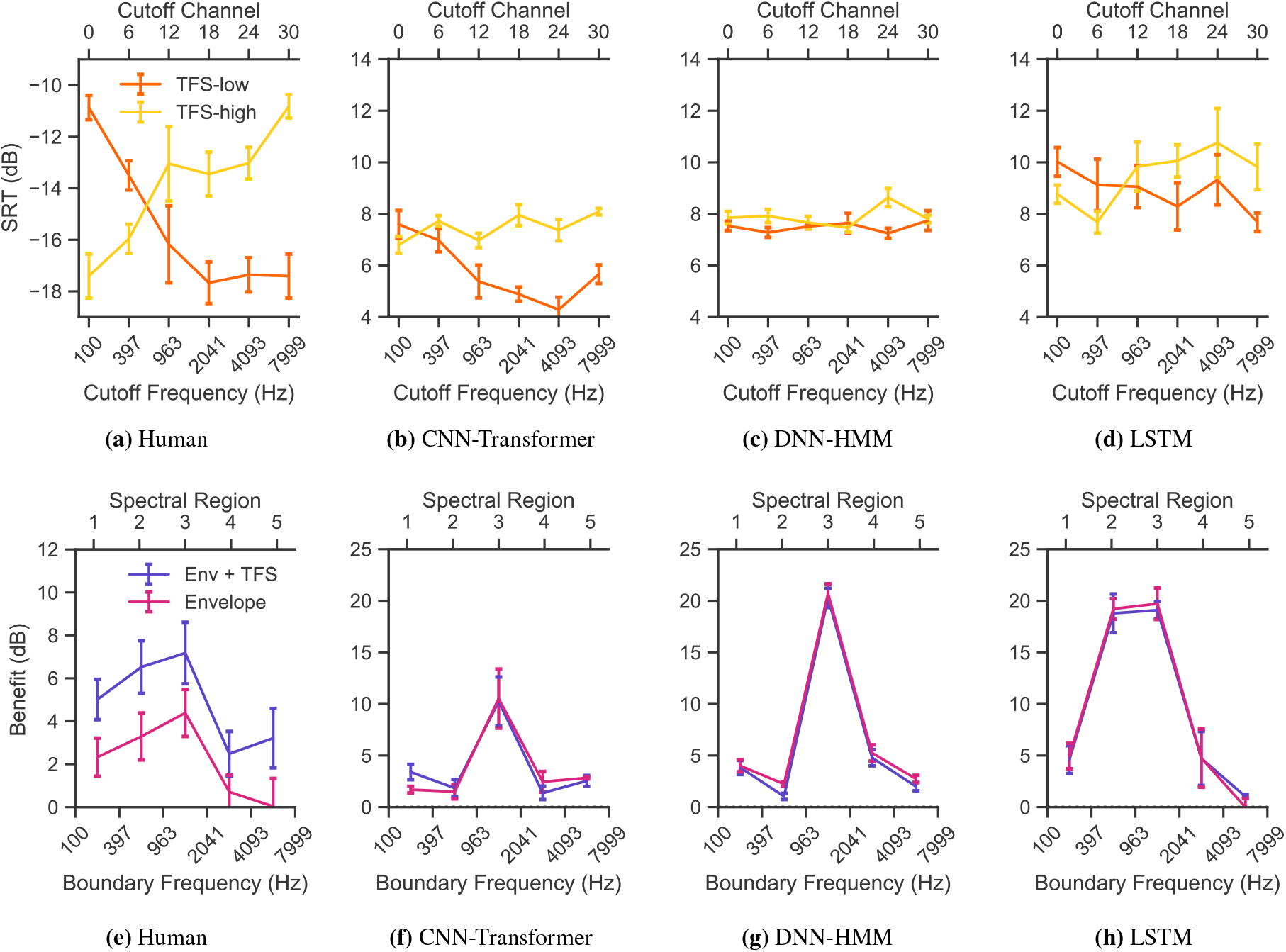
Same as in Figure 3 but here performance is measured against 100 sentences from the CRM dataset (single male talker) rather than IEEE sentences

